# Lymphoid tissues contribute to viral clonotypes present in plasma at early post-ATI in SIV-infected rhesus macaques

**DOI:** 10.1101/2023.05.30.542512

**Authors:** Antonio Solis-Leal, Nongthombam Boby, Suvadip Mallick, Yilun Cheng, Fei Wu, Grey De La Torre, Jason Dufour, Xavier Alvarez, Vinay Shivanna, Yaozhong Liu, Christine M. Fennessey, Jeffrey D. Lifson, Qingsheng Li, Brandon F. Keele, Binhua Ling

## Abstract

The rebound-competent viral reservoir (RCVR), comprised of virus that is able to persist during antiretroviral therapy (ART) and mediate reactivation of systemic viral replication and rebound viremia after antiretroviral therapy interruption (ATI), remains the biggest obstacle to the eradication of HIV infection. A better understanding of the cellular and tissue origins and the dynamics of viral populations that initiate rebound upon ATI could help develop targeted therapeutic strategies for reducing the RCVR. In this study, barcoded SIVmac239M was used to infect rhesus macaques to enable monitoring of viral barcode clonotypes contributing to virus detectable in plasma after ATI. Blood, lymphoid tissues (LTs, spleen, mesenteric and inguinal lymph nodes), and non-lymphoid tissues (NLTs, colon, ileum, lung, liver, and brain) were analyzed using viral barcode sequencing, intact proviral DNA assay, single-cell RNA sequencing, and combined CODEX/RNAscope/*in situ* hybridization. Four of seven animals had viral barcodes detectable by deep sequencing of plasma at necropsy although plasma viral RNA remained < 22 copies/mL. Among the tissues studied, mesenteric and inguinal lymph nodes, and spleen contained viral barcodes detected in plasma, and trended to have higher cell-associated viral loads, higher intact provirus levels, and greater diversity of viral barcodes. CD4+ T cells were the main cell type harboring viral RNA (vRNA) after ATI. Further, T cell zones in LTs showed higher vRNA levels than B cell zones for most animals. These findings are consistent with LTs contributing to virus present in plasma early after ATI.

**One Sentence Summary:** The reemerging of SIV clonotypes at early post-ATI are likely from the secondary lymphoid tissues.

## Introduction

Persistence of the rebound-competent viral reservoir (RCVR), even with extended time on ART, represents the primary obstacle to achieving a cure of HIV infection. All currently approved antiretroviral treatments can block new rounds of infection but do not impact provirus in latently infected cells. Once ART blockade of new infections is stopped, ongoing proviral reactivation can lead to viral spread and rebound viremia. If not re-suppressed, this reignited productive infection will eventually lead to progressive immunodeficiency and AIDS, along with a risk of viral transmission (*1*). A better understanding of the sources, timing, and mechanisms of viral rebound post-antiretroviral therapy interruption (ATI) should help to develop strategies to target and reduce or eliminate these reservoirs (*1*).

The RCVR may include any cells containing an intact provirus capable of reactivation leading to systemic productive infection. It is possible that non-lymphoid tissues (NLTs) or organs, such as the gastrointestinal tract, genital tract, adipose tissue, lung, and brain, (*2, 3*), which contain fewer latently infected cells than lymphoid tissues (LTs), may rekindle HIV productive infection. However, LTs, which contain abundant CD4+ T lymphocytes, have proven to be a major tissue reservoir during suppressive ART (*4, 5*). Human LTs are highly diverse and complex in structure, function, and anatomic location. There are approximately 800 lymph nodes (LN) distributed throughout the body in humans (*6*). We hypothesize that LTs, harboring a greater number of replication-competent proviruses, in proximity to available CD4+ T cells for local viral spread and replication during ATI, are the predominant sources of viral rebound. While identifying anatomic sources of viral rebound in people living with HIV could be informative, such analyses are impractical due to the challenges in sampling relevant tissue sites. Most studies assessing rebounding virus during ATI only sampled peripheral blood. However, the rebounding virus population in blood may reflect a mixed contribution from many distinct tissue sites. Thus, the tissue origins of the initial rebounding virus during early ATI remain largely unknown and a comprehensive analysis of all potential tissue sources of HIV rebounding viruses is needed.

To fill the gap and attempt to identify potential sources of rebounding virus during early ATI, we employed a rhesus macaque model that permitted extensive tissue sampling. We infected macaques with the barcoded virus, SIVmac239M (*7, 8*). SIVmac239M is a modified SIVmac239 with genetically introduced barcodes consisting of 34 randomized nucleotide bases inserted between the *vpx* and *vpr* genes in an otherwise genetically-identical backbone. This viral stock contains over 10,000 distinct viral barcodes which can be used to track individual viral clonotypes over time in blood and tissues (*7*). Importantly, barcode variants have comparable replicative capacity and the barcodes remain intact during a long replication period and have been used to quantify and characterize rebounding viral clonotypes in plasma following ATI (*7, 9*). Herein, we compared viral barcodes detectable at necropsy in plasma (SIV RNA < 22 copies/mL), various LTs, and NLTs by deep sequencing in early post-ATI to identify shared barcodes in plasma and tissues to infer potential sources of rebound virus.

After infection, seven Chinese-origin rhesus macaques underwent two rounds of ART and ATI. ATI-1 was intended to evaluate the timing and extent of viral rebound in plasma, the composition of viral clonotypes in peripheral blood after long-term ART, and to examine the potential influence of a prior ATI on viral barcodes detected after a subsequent ATI. One week after the second ATI (post-ATI-2), when all macaques still had plasma viral RNA levels < 22 copies/mL, necropsies were performed and multiple LTs (mesenteric LN (MesLN), inguinal LN (IngLN), and spleen), NLTs (ileum, colon, lung, liver, and brain) and blood from necropsy, were examined by viral barcode sequencing, cell-associated DNA (CA-DNA) and cell-associated RNA (CA-RNA), intact proviral DNA assay (IPDA), combined CODEX/RNAscope/*in situ* hybridization, single-cell RNA sequencing (scRNAseq), and other approaches. The results suggested that among the tissues studied, MesLN, IngLN, and spleen showed overlap with viral barcodes detectable by sensitive sequencing of plasma at necropsy, suggesting that these tissues may contribute to viral reactivation and eventual rebound, post-ATI.

## Results

### Dynamics of longitudinal plasma viral load during viral infection, ART, and ATI

Plasma viral load (pVL) from peripheral blood of infected rhesus macaques was assessed longitudinally (**Fig. 1**). Viremia peaked at around 2 weeks post-infection (wpi), ranging from 6.3x10^6^ to 2.7x10^7^ copies/mL. When ART was initiated at 12 wpi, the animals had reached a set point with pVLs ranging between 1.2x10^3^ to 2.6x10^4^ copies/mL. Chronic pVL values in the Chinese rhesus macaques in the present study were relatively lower than previous reports of SIVmac239M infection in Indian rhesus macaques, which is consistent with past comparisons of SIVmac lineage viruses in Indian vs. Chinese-origin rhesus macaques (*10*). During the 42 weeks of daily administration of ART, minor viral blips were detected in KN41, KT12, LB66, and LC31 at 20 wpi, in DJ01 and LB66 at 24 wpi, in DJ01 and KT12 at 28 wpi, in LB66 at 36 wpi, and EM77 at 40 wpi, but otherwise, pVL was below the assay limit of detection of 81 copies/mL. Selected time points (28, 32, 36, and 40 wpi) during ART as well as during ATI-2 were further evaluated by a more sensitive assay and were found to be less than the limit of detection (22 copies/mL). After the first ART interruption (ATI-1) at 52 wpi, the plasma vRNA values at 54 wpi ranged from < 22 to 10^3^ copies/mL. By 57 wpi, plasma vRNA values had increased to from 3.47x10^2^ to 2x10^4^ copies/mL. After 3 weeks off treatment, ART was resumed and pVL quickly dropped below 81 copies/mL and remained under 22 copies/mL until euthanasia and necropsy (68-75 wpi), which occurred one week following the second ART interruption (ATI-2).

**Fig. 1:**
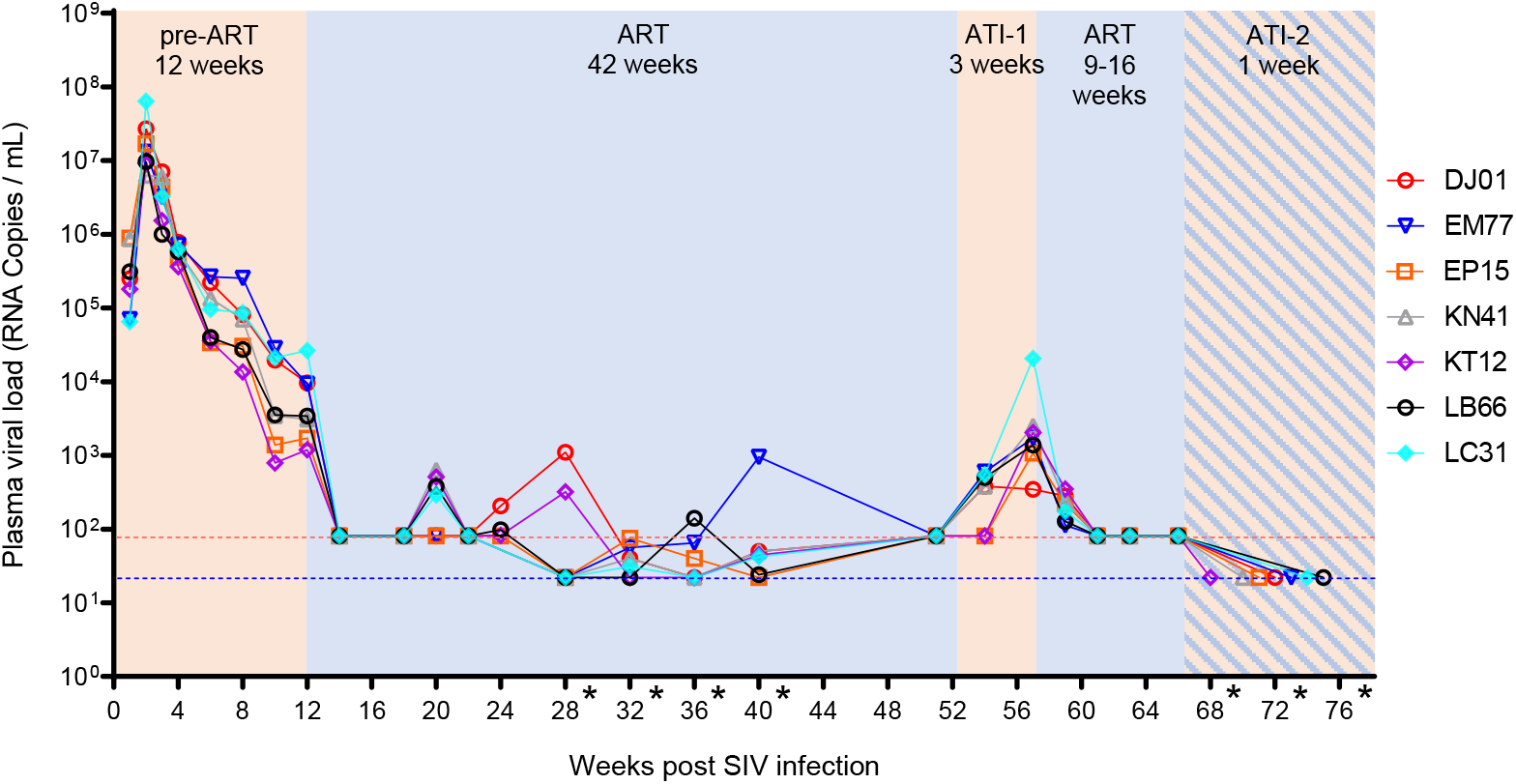
Experimental design and dynamics of SIV plasma viral load (pVL). The limit of detection of this assay was 81 copies/mL (red dashed line), and the samples from 28, 32, 36, and 40 wpi, and necropsy were further tested by a more sensitive Q-RT-PCRT with a limit of detection of 22 copies/mL (blue dashed line). These time points were marked with an asterisk (*). pVL was undetectable at the necropsy timepoint during ATI-2.

### Dynamics of viral clonotypes in blood and detection of barcodes in plasma at one week of ATI-2

Barcode sequencing was performed on plasma samples during acute infection (week 2), chronic infection/pre-ART (week 12), ATI-1 (week 57), and ATI-2 (weeks 68-75) (**Fig. 2**). As expected, given the high administered dose of SIVmac239M, a wide variety of barcodes were detected in the peak and chronic phases. However, the number of clonotypes detected decreased considerably in the 10 weeks from peak to chronic phase and between pre-ART and both ATI time points. At peak viremia, there were an average of 741 detectable barcodes (range: 388-1519), which decreased during chronic viremia to only an average of 77 (range:10-217), likely reflecting the effects of early viral dynamics and stochastic differential selective processes operating on otherwise initially identical viral variants bearing distinct viral barcodes. At ATI-1, barcode sequencing revealed an average of 8.3 clonotypes in the plasma viral population (range: 1-19). These rebounding viral lineages were most often but not exclusively found within the top 10% of the peak or chronic phase barcode distribution (**Fig. 2**).

**Fig. 2:**
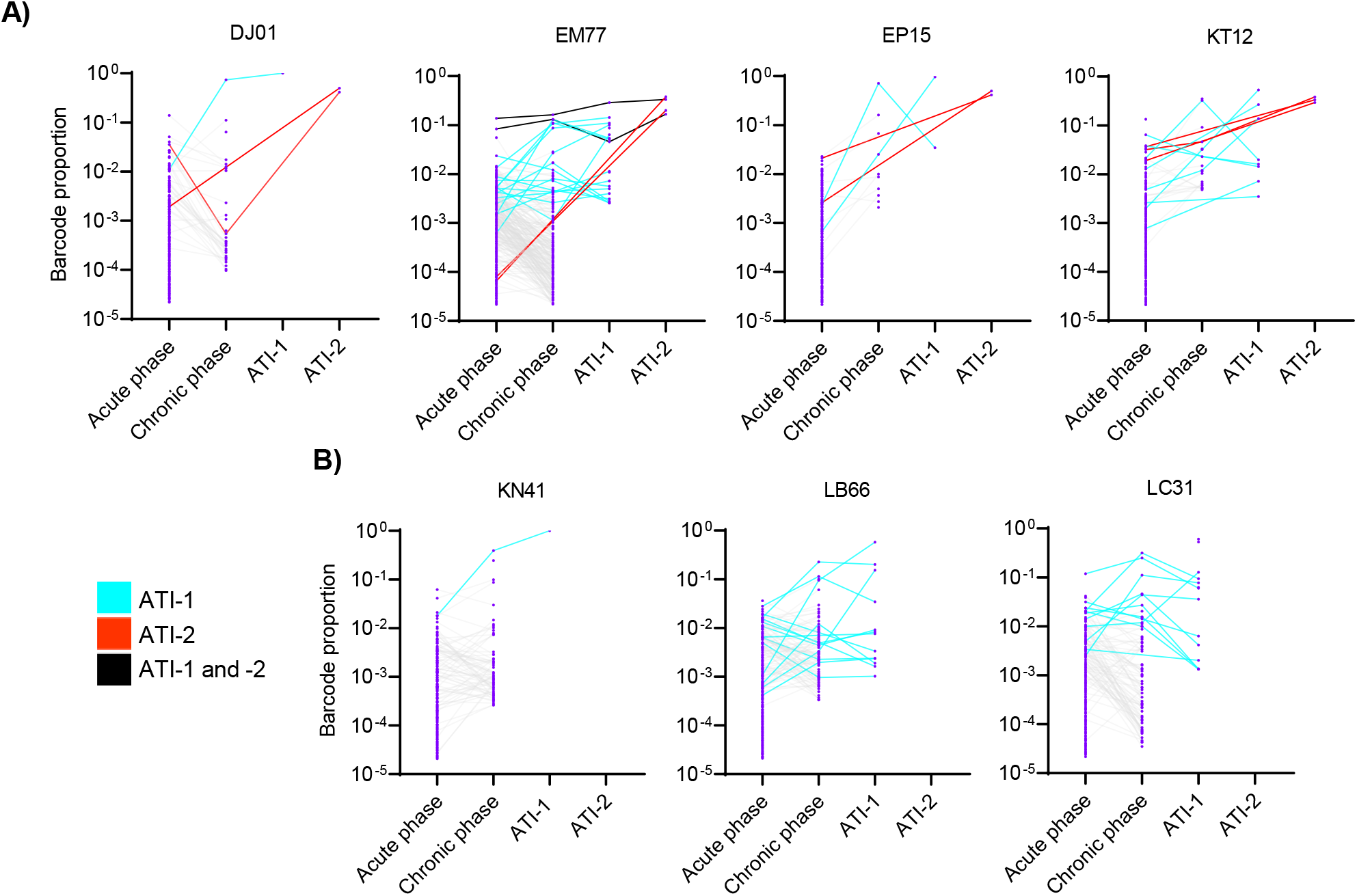
Detection of SIVmac239M barcodes in plasma during acute and chronic phases of SIV infection, ART, and ATIs. **A)** Animals in which barcodes were detectable in plasma during ATI-2 (Group 1). **B)** Animals in which no viral barcodes were detected in plasma during ATI-2 (Group 2).

During ATI-2, although plasma SIV RNA was below 22 copies/mL for all animals, more sensitive sequencing techniques detected viral barcode sequences in plasma at necropsy from 4 of 7 animals (DJ01, EM77, EP15, KT12) (group 1). Viral barcodes detected in plasma at ATI-2 were often not the dominant pre-ART barcodes and in some cases were derived from minor lineages (EM77). The remaining three animals (KN41, LB66, and LC31) had no detectable viral barcodes in plasma at ATI-2 (group 2).

In addition, viral barcode sequences were also found in CA-DNA from peripheral blood mononuclear cells (PBMCs), with barcodes detected during ATI-1 and ATI-2 also identified in PBMCs from acute and chronic phases (Fig. S1). Similar to plasma, a variety of clonotypes were detected in all of the subjects at different time points.

### Viral clonotype diversity and levels of intact proviruses in PBMCs

The frequency of viral barcode clonotypes found in plasma during ATI-1 that were also found in matching PBMC was larger in group 1 animals compared to group 2. Although this difference was not statistically significant, it was consistent for all of the time points studied (**Fig. 3A**). In addition, the size of the intact proviral reservoir in PBMCs, as inferred by IPDA, showed a similar pattern of being nominally larger throughout the study in animals from group 1 (**Fig. 3B**). Furthermore, the reduction of the viral barcode clonotype diversity over time, previously observed in plasma, was also manifested in CA-DNA from PBMCs.

**Fig. 3:**
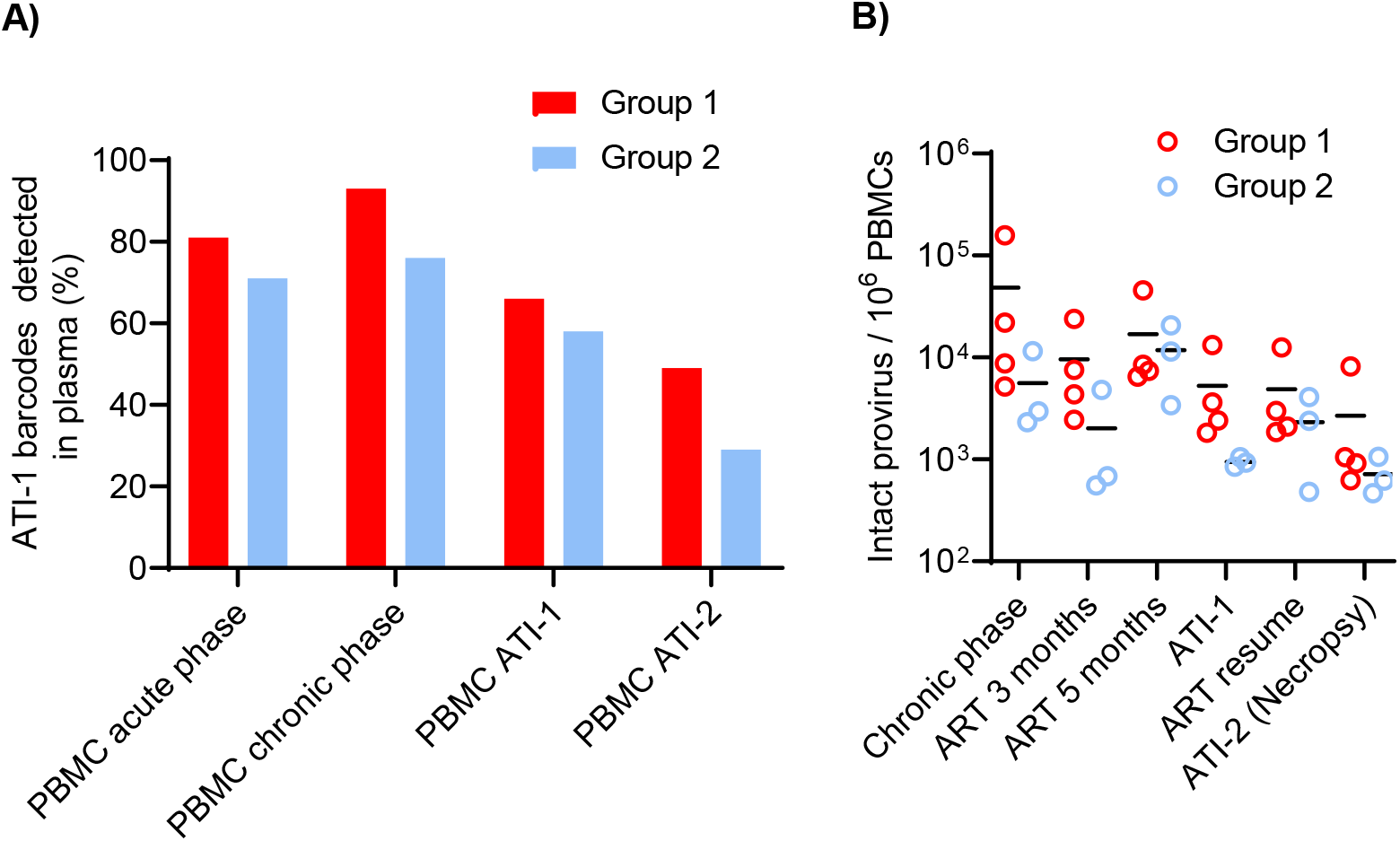
Longitudinal assessment of barcode clonotype numbers and the size of viral reservoir in PBMCs. **A)** Normalized detection of SIVmac239M barcodes in DNA from PBMCs at different stages that supported the viral rebound at ATI-1 in plasma in at least 1% of the total barcodes detected. **B**) Levels of viral reservoir in PBMCs at multiple time points in both group of animals .

### LTs contained more clonotypes than all the other tested tissues at ATI-2

Viral loads of CA-DNA and CA-RNA from blood, LTs (MesLN, IngLN, spleen), and NLT (colon, ileum, lung, liver, and brain) were quantified. Most of these tissues showed measurable vRNA and/or DNA levels (**Fig. S2**). Next, we assessed the total number of different clonotypes present and expressed in each type of tissue from all the animals (**Fig. 4**). Interestingly, the highest mean number of distinct clonotypes was found in the MesLN (44.0±17.0, mean±SD) and IngLN (39.3±33.8), while lower numbers of clonotypes were detected in the spleen (11.3±5.6), colon (3.9±7.0), ileum (3.1±7.5), lung (1.1±1.7), and liver (0.1±0.4, **Fig. 4A**). Sequence analysis of complementary DNA produced from CA-RNA revealed high levels of distinct viral barcodes in CA-RNA from MesLN and IngLN, as seen for viral DNA. The MesLN (8.7±6.1, mean±SD) contributed the highest frequency of the total resident clonotypes, followed by the spleen (6.1±4.7), IngLN (5.1±7.2), colon (1.7±3.4), lung (1.0±1.3), and basal ganglia of the brain (0.1±0.4, **Fig. 4B**). Meanwhile, the number of detected clonotypes in viral DNA was positively correlated with the number of detected clonotypes in vRNA in these tissues (**Fig. 4C**).

**Fig. 4:**
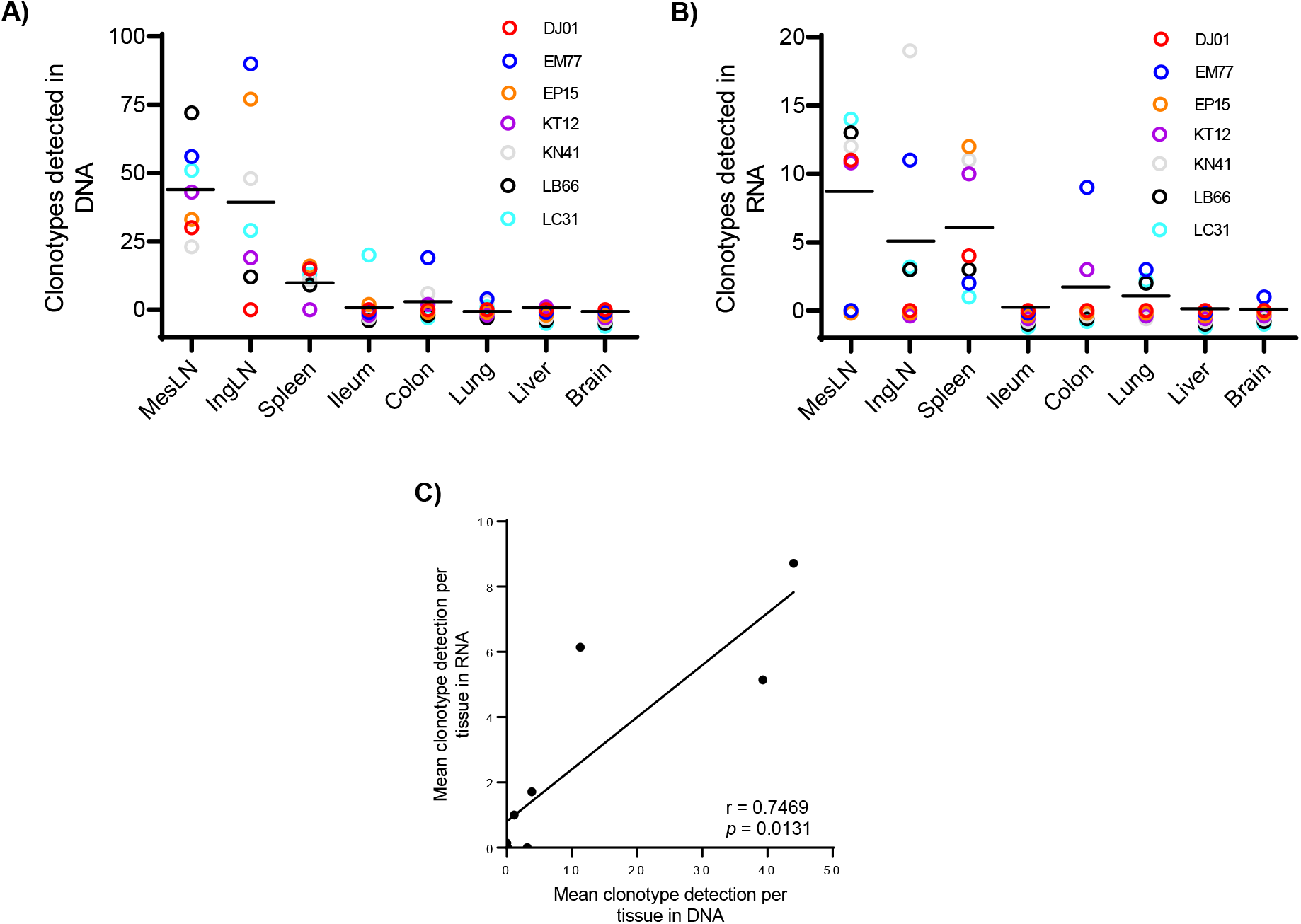
Tissue viral clonotype numbers from CA-DNA or CA-RNA at ATI-2. **A)** Number of clonotypes in each tissue from CA-DNA samples. **B)** Number of clonotypes in each tissue from CA-RNA samples. **C)** CA-DNA mean values for each tissue paired with their respective CA-RNA mean value showed a statistically significant positive correlation (*p*=0.0131).

### Presence of shared viral barcodes in tissues and ATI plasma

To assess the relative presence of each clonotype in each tissue (barcode proportion) and compare to the barcodes detected in plasma at ATI-1 and ATI-2 for animals in group 1, we sequenced viral DNA and identified barcodes from multiple tissues (**Fig. 5** and **Fig. S3)**. In general, the clonotypes detected in ATI-2 plasma were better represented in DNA from necropsy MesLN, IngLN, and spleen, compared to the other tissues studied. (**Fig. 5C**).

**Fig. 5:**
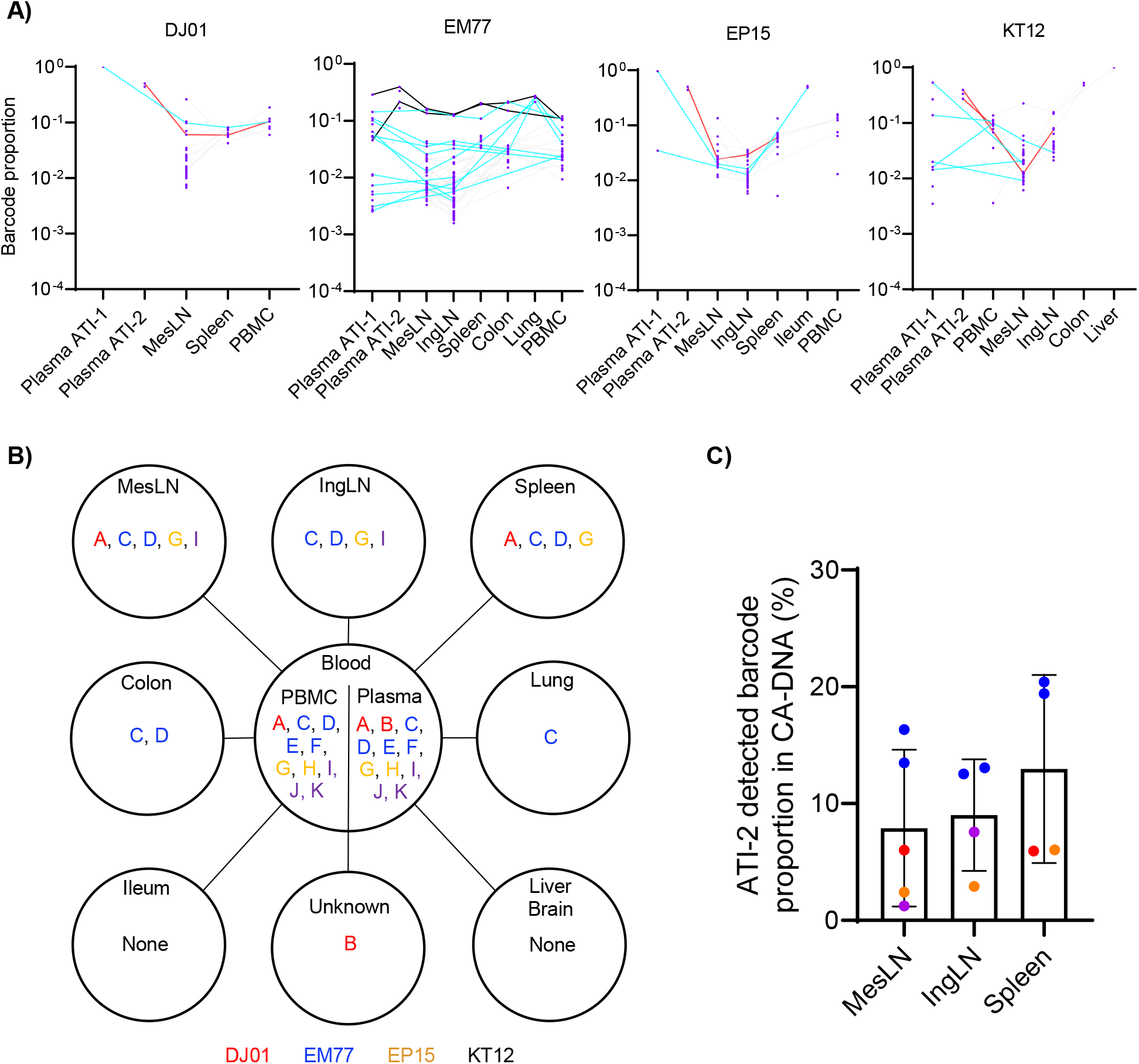
SIVmac239M barcode detection in CA-DNA from 9 tissues corresponding with plasma barcode detection at ATI-1 and ATI-2 in group 1. **A)** Clonotypes found in each animal. The barcodes from plasma viral RNA at ATI-1 and ATI-2 were included as reference related to rebound. Barcodes associated with ATI-1 are shown with a cyan line. Barcodes associated with ATI-2 are shown with a red line. Barcodes associated with ATI-1 and ATI2 are shown with a black line. **B)** The eleven barcodes detected in plasma during ATI-2 are represented by the letters A to K and the viral clonotypes are divided by the animals in which they were detected as follows: DJ01 (red, A-B), EM77 (blue, C-F), EP15 (gold, G-H) and KT12 (purple, I-K). **C)** Tissue distribution of the total proportion of the ATI-2 detected plasma viral clonotypes.

Meanwhile, despite detection of fewer clonotypes in plasma compared to CA-DNA, the analysis of viral barcodes obtained from CA-RNA found that the clonotypes detected in ATI-2 plasma were also more frequently present in the MesLN, IngLN, and spleen at a higher proportion than the other tissues studied (**Fig. S4 and S5**).

Of note, although no virus was detected in plasma in animal KN41 of group 2 during ATI-2 either by Q-RT-PCR or by barcode analysis, some viral barcodes were found in plasma at 68 wpi just prior to necropsy. There was overlap between the barcodes from this timepoint and barcodes detected at necropsy in CA-DNA from MesLN, IngLN, colon, and lung. (**Fig. S6)**.

### Higher levels of intact proviruses were found in MesLN and spleen

Given that LTs harbored more clonotypes that overlapped with those detected in ATI-2 plasma, we next assessed the size of inferred intact proviral population in MesLN and spleen. CD4+ T cells from both tissues showed high levels of intact provirus in all of the animals, with group 1 having a significantly higher level compared to group 2 (**Fig. 6A**). CD11b+ myeloid cells from MesLN and spleen tissues also contained intact proviruses (**Fig. 6B**) but these were only found in one or two animals, respectively.

**Fig. 6:**
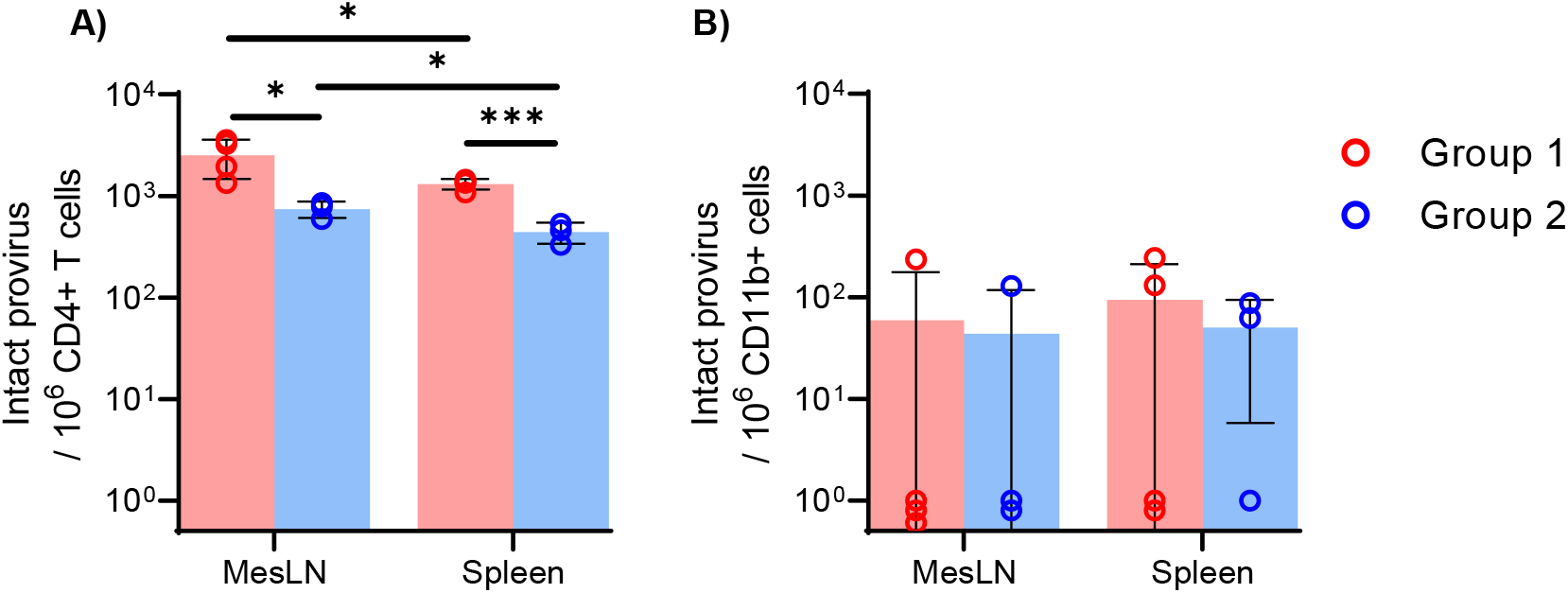
Levels of SIV intact provirus in tissues at ATI-2. **A)** Intact provirus per 1x10^6^ CD4^+^ T cells in MesLN and spleen. **B)** Intact provirus in CD11b+ cells in MesLN and spleen. * *p* < 0.05, and *** *p* < 0.001.

### Tissue vRNA was mainly detected in CD4+ T cells from MesLN and spleen

While multiple studies have already shown that CD4+ T cells are major cellular viral reservoirs in lymphoid tissues during ART (*4, 11–13*), we wanted to determine the cell types actively producing vRNA in these sites at the early phase of ATI (ATI-2) prior to detectable systemic spreading. Using SIV RNAscope, we found that CD4+ T cells were the main cell type expressing vRNA (**Fig. 7A, Fig. S7A, and S7B**) while CD11b+ myeloid cells produced limited vRNA signal (**Fig. S7C-S7E**). We then quantified vRNA in both MesLN and spleen in all 7 animals (**Fig. S8A**). When separately comparing vRNA levels in MesLN and spleen between group 1 and group 2, there were no significant differences (**Fig. S8B and S8C).** However, it is noteworthy that MesLN showed a significantly higher number of vRNA-positive cells compared to the spleen across the 7 animals (**Fig. 7B**). This result agrees with the findings of a larger intact proviral reservoir (**Fig. 6A**) and a higher number of distinct clonotypes in MesLN than in spleen (**Fig. 4**).

**Fig. 7:**
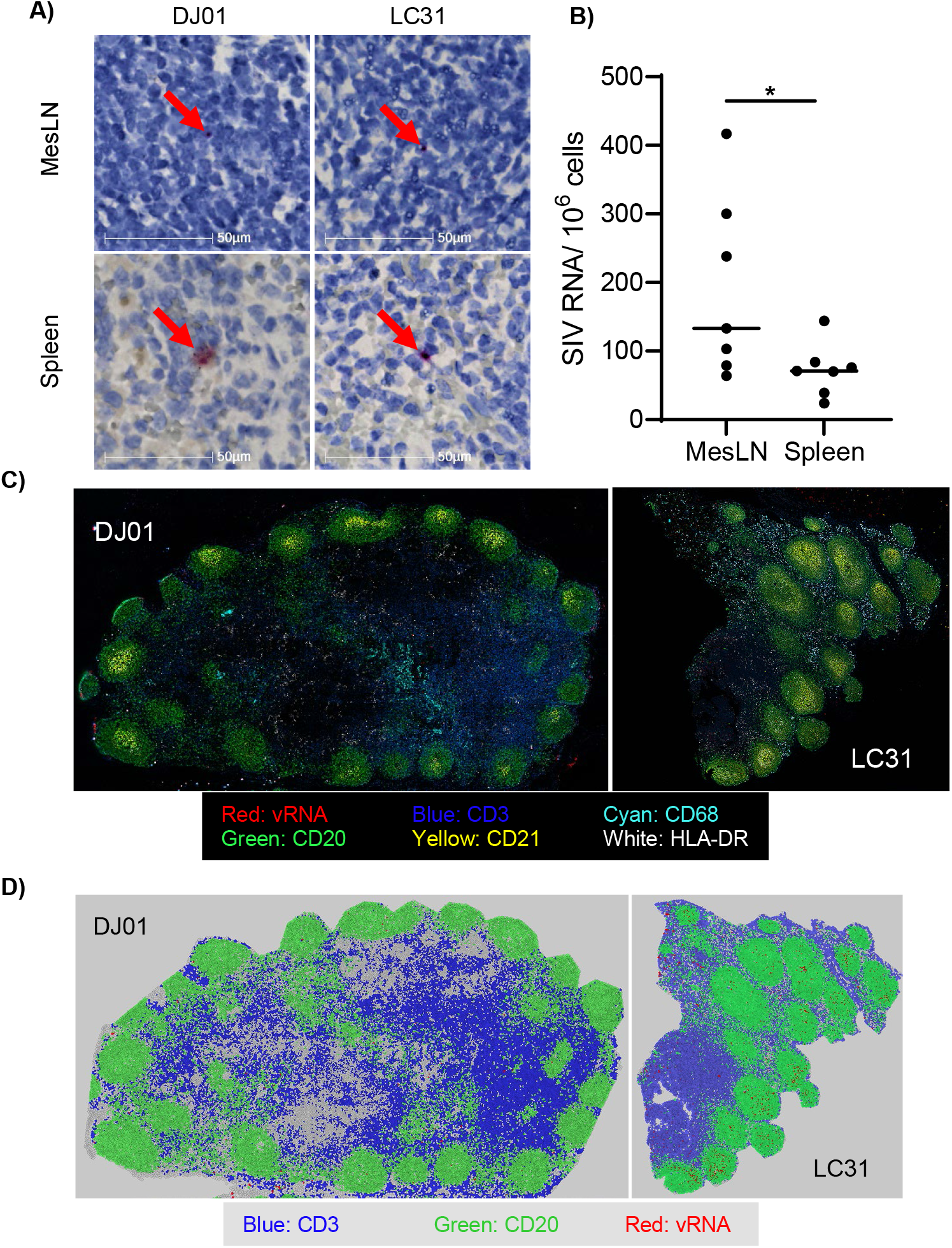
Distribution of detected SIV RNA-positive cells in MesLN and spleen at ATI-2. Slides were fully scanned to detect and quantify the presence of vRNA in cells. **A)** Representative vRNA detection in MesLN and spleen from animal DJ01 in group 1 and animal LC31 in group 2. **B)** Comparison of levels of vRNA-positive cells from all the animals between MesLN and spleen. *: *p* < 0.05. **C)** Com-CODEX-RNAscope image of MesLN from animals DJ01 and LC31 **D)** Voronoi plot of the Com-CODEX-RNAscope shows the vRNA (red) distribution in the B cell zones (marked by CD20 in green) and T cell zones (Marked by CD3 in blue).

We then applied the combined CO-detection by indexing (CODEX) and RNAscope *in situ* hybridization (CODEX-RNAscope) approach for characterizing spatial relationships between immune cells and virus expression. Similar to our previous findings on animals under ART (*11*), we readily detected vRNA in MesLNs of both groups (**Fig. 7C and 7D**, and **Fig. S9**). Here, we further identified the spatial distribution of vRNA-positive cells in T and B cell zones. T cell zones showed a higher vRNA level than B cell zones for most animals with the log add ratio of T cell *vs* B cell zones of DJ01, 0.05 *vs* 0.01, EM77, 0.54 *vs* 0.12, LB66 0.73 *vs* 0.1, and LC31 0.48 *vs* -0.26. Interestingly, a different distribution pattern of vRNA expression was observed between groups. While the vRNA signals of group 2 tended to be clustered, consistent with localized viral expression, the vRNA signals of group 1 were already spread more evenly across the tissue section.

### Multiple genes in MesLNs were dysregulated in animals with viral barcodes detected in plasma at ATI-2

To understand potential cellular mechanisms that differentiate animals with and without detectable viral barcodes in plasma at ATI-2, the cells isolated from MesLNs extracted at necropsy from different animals of each group were analyzed by single-cell RNA sequencing (scRNA-seq) and transcriptomic analysis. Five cell clusters were initially differentiated, of which 4 were annotated, showing markers of B cells, Naïve T cells, CD4+, and CD8+ T cell subsets. These clusters had similar proportions between the two groups, with small differences, especially in the proportion of Naïve T cells (**Fig. 8A and 8B, Table S1**). Resolution, which is a parameter of the Louvain detection algorithm that modify the number of recovered clusters, was increased in the data processing module of the software from a low-resolution (0.1), to a high-resolution (2.0). This option sets the granularity of the downstream clustering, with increased values leading to an enhanced cluster differentiation. These clusters were further annotated into multiple cell subsets within the initial cell clusters (**Fig. S10A**). Again, all the clusters were found in both groups allowing for comparison (**Fig. S10B**).

**Fig. 8.**
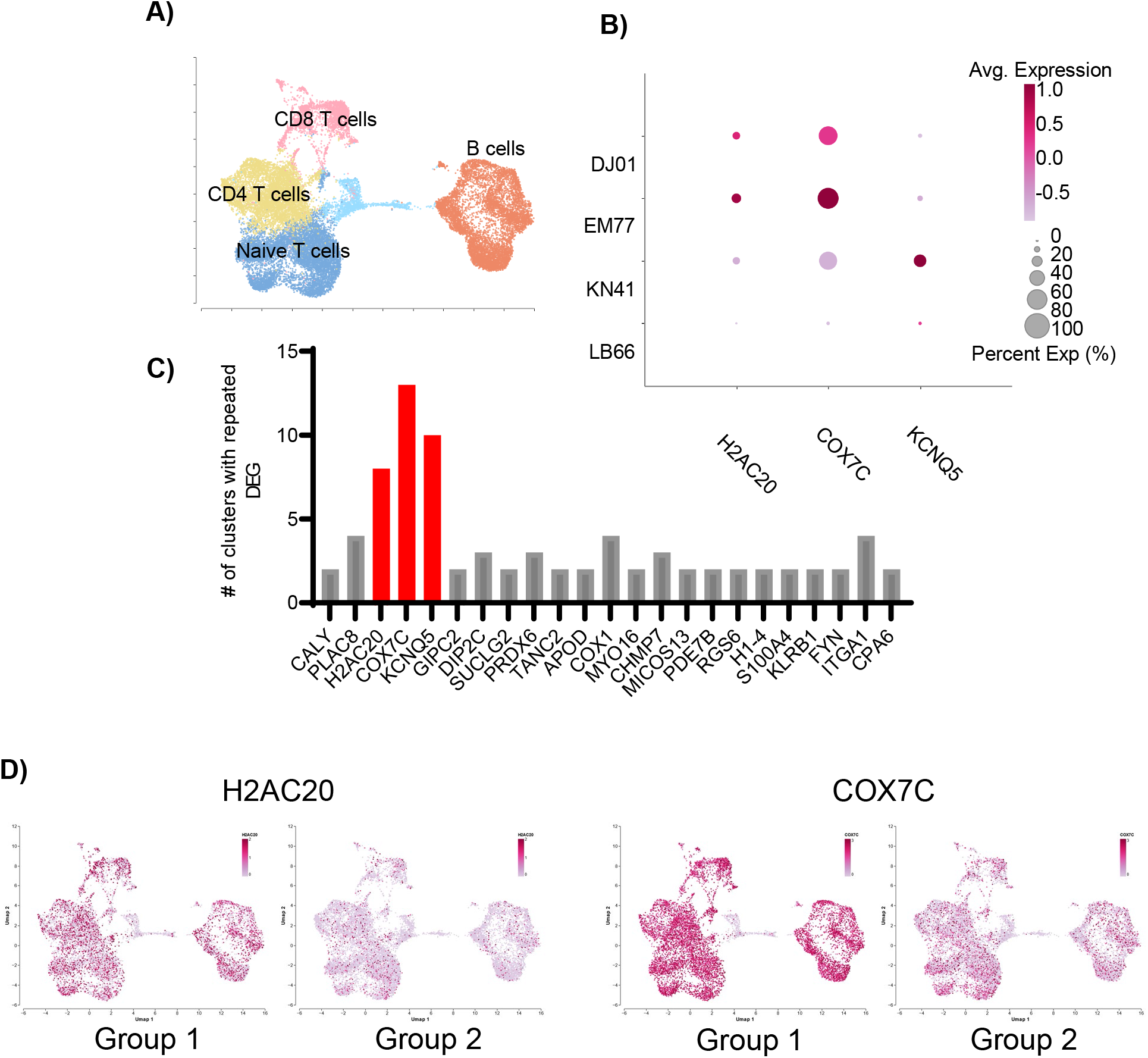
scRNAseq analysis of MesLN isolated cells at ATI-2. **A)** Louvin clusters at 0.1 resolution. **B)** Dot plot of the genes frequently dysregulated across multiple louvin clusters in group 1 animals. **C)** DEGs repeated across different louvin clusters at 2.0 resolution. **D)** Differential expression of H2AC20 and COX7C in the two groups.

A total of 91 differentially expressed genes (DEGs) were found across all the cell subsets, of which, multiple were related to an enhanced immune activation (**Table S2**). The highest logarithmic fold change (LogFC) in all the subsets between group 1 and group 2 was observed in follicular helper CD4+ T cells in the gene *SPINK2* (LogFC:4.369, adj *p*-value: 0.037), and *AFDN* (LogFC:2.5403, adj *p*-value:0.045, **Table S3**). Interestingly, from all the DEGs listed in each cell subset analyzed **(Table S3),** approximately 24 genes were identified that were similarly dysregulated among different cell clusters. Genes such as *COX7C, H2AC20*, and *KCNQ5* were modulated in 8∼13 clusters including multiple B and T cell subsets under the 2.0 resolution analysis (**Fig. 8C-8E**). Specifically, *COX7C,* which is related to mitochondria dysfunction, increased expression in 13 clusters, together with other genes related to mitochondrial processes and oxidative stress that were moderately dysregulated. While *KCNQ5* was downregulated in 10 clusters, other genes related to its function were not changed. *H2AC20*, which is involved in nucleosome and histone modifications, was upregulated in 8 clusters together with other dysregulated genes in this category (**Table S2**).

## Discussion

A better understanding of the origin of the virus that initiates viral rebound upon ATI may help develop better therapeutic strategies to minimize or eradicate the RCVR to facilitate a cure for HIV infection. While Rothenberg et al. documented multifocal viral recrudescence in LTs following ATI (*14*), it remains unknown whether there are preferred sites of early viral reactivation following ATI, and thus identifying such sites may be useful in the development of effective interventions. Since there are several hundred lymph nodes that may contribute to the RCVR, many of which are found in distinct anatomic locations and are associated with different microenvironments that may potentially result in different immune activation states, a survey of many LTs and NLTs potentially contributing to early viral reactivation and rebound is needed. While not definitive due to limitations including sampling timing and the number and amounts of tissues sampled, our study nonetheless explores the possibility that the source of virus rebound might be identified at a time right before systemic viral spreading is measurable. Through comparative analysis of virus clonotypes detectable in plasma and multiple LTs and NLTs, at necropsy, one week after ATI-2, we found that the viral barcodes detected in plasma at necropsy were also found in MesLN, IngLN, and spleen, consistent with the possibility that these tissues may contribute to post-ATI viral reactivation and eventual rebound.

Our data also showed a large pool of intact sequences and potentially replication-competent proviruses in these LTs as inferred from IPDA analysis. The availability of CD4+ T cells in these tissues enables the local spread of reactivated virus expression and render distinctive features of tissue microenvironments. For instance, the MesLNs drain the contents from the gut and are more immunologically activated than many other lymphoid tissues, particularly in HIV infection (*15*). MesLN exhibited a larger intact proviral pool and higher levels of vRNA expression than the spleen. Interestingly, despite the larger number of distinct viral clonotypic barcodes in MesLN, spleen, and IngLN compared to PBMCs, most of the barcodes found in plasma at ATI-2 were also identifiable at some point in CA-DNA from PBMCs. It is not possible to determine if these plasma viral clonotypes originated in PMBCs (*16–18*) or if these lymphocytes simply represent the trafficking of lymphocytes from various LTs. In addition, we cannot exclude the possibility that the viral clonotypes detected in plasma during ATI-2 could have been produced by a low residual expression during the ART period, and not solely by emerging proviral reactivation.

In addition, IngLN also showed distinct viral clonotypes that were also present in the ATI-2 plasma and had high diversity in CA-DNA and CA-RNA, suggesting that IngLN might have been underappreciated as a site of SIV persistence during ART and as a potential source of viral recrudescence (*4*). Given that it is a relatively easier site to access than MesLN, it may be informative to sample IngLN in clinical studies involving people living with HIV.

Despite previous observations regarding the capacity of the small intestine to contain large SIV pools (*19, 20*), no viral barcodes detected in ATI-2 plasma were found in the samples of small intestine herein analyzed, and only one animal from group 2 showed detectable intact proviral DNA there. Similarly, only one animal had a viral barcode clonotype detected in the colon sample that was also found in plasma during ATI-2. When interpreting this finding, it is important to recognize that we were only able to sample and analyze a very small portion of the total gastrointestinal tract, which despite containing a major fraction of the total lymphoid cells in the body, has these cells interspersed discontinuously amidst a dominant population of non-lymphoid cells, posing sampling challenges. More extensive sampling and analysis of each tissue could potentially add to the current evidence suggesting a primary role for secondary lymphoid tissues contributing to early viral rebound.

The viral, cellular, and tissue factors that facilitate viral reactivation and the associated mechanisms that underlie viral recrudescence in lymphoid tissues, such as the MesLN, remain to be further explored. In this study, our single-cell RNA transcriptomic analysis of MesLN revealed differences between the two groups, including dysregulation of genes related to mitochondria function and oxidative stress. Dysregulation of mitochondria associated with HIV/SIV infection has been previously associated with the disruption of ATP synthesis, increased oxidative stress, and the release of cytochrome c, resulting in HIV-induced apoptosis (*21, 22*). On the other hand, the observed dysregulation of genes related to the nucleosome and histone modifications may impact cellular epigenetics, potentially affecting the binding of transcription factors or the cell transcriptome (*23, 24*). The chromatin remodeling induced by HIV has been previously reported to have the potential to affect several cellular pathways and promote latency in viral reservoirs (*25–28*). The dysregulation of this mechanism in the MesLN could be further studied to elucidate whether it could be one potential cause of the formation of a larger SIV reservoir. Interestingly, our results found that the *KCNQ5* gene was downregulated in animals with plasma barcode detection at ATI-2. This is a member of the KCNQ potassium channel gene family and despite scarce reports suggesting a link between KCNQ5 modulation and HIV pathogenesis, more research is needed to determine if an association with any cellular pathways after HIV/SIV infection really exists (*29–32*).

Reduction or elimination of the RCVR for HIV or SIV has proven to be very challenging for many reasons, including the significant differences between the organs, tissues, and cells containing intact proviruses. LTs, such as lymph nodes and spleen, harbor the majority of CD4+ T cells and are the major sites of HIV persistence during ART (*12, 33, 34*). In these tissues, subsets of CD4+ T cells harboring replication-competent virus include central, transitional, and effector memory T cells, tissue-resident follicular helper T cells, and possibly naïve T cells (*2, 35–37*). Likewise, myeloid cells can be infected, and specifically, peripheral monocytes and macrophages (*38–40*) have shown resistance to HIV-induced apoptosis (*41*). Similarly, microglial cells and perivascular macrophages can contribute to viral persistence in the central nervous system (CNS) (*2, 3*). Viral rebound after ART discontinuation is thought to be produced by a small number of cells in different tissues, creating a multifocal origin of the rebound (*33, 42, 43*). More efforts aimed at better understanding the sources of early viral reactivation and rebound after ATI in animal models and in humans could aid in the development of therapies to avoid it.

In conclusion, our results suggest that LTs, especially MesLNs, may be important contributors to the very early virus detectable in plasma post-ATI, even when plasma vRNA is < 22 copies/mL. However, further studies are needed to evaluate the relative potential contributions from other tissues and organs. Our results also suggested that despite the wide distribution of replication-competent proviruses across the body observed in both animals and humans (*1, 4, 44*), a limited number of lineages from one or multiple tissues may contribute to the earliest viral population detectable in plasma once suppressive ART is discontinued.

## Methods

### Experimental design

Animals were housed at the Tulane National Primate Research Center and maintained following the standards of the American Association for Accreditation of Laboratory Animal Care, the “Guide for the Care and Use of Laboratory Animals” prepared by the National Research Council, and the Animal Welfare Act. All studies were approved by the Tulane Institutional Animal Care and Use Committee (IACUC). Seven rhesus macaques (Macaca mulatta) of Chinese origin were studied. Animals were sero-negative for SIV, simian D retrovirus, and simian T-cell leukemia virus before SIV inoculation. All of them underwent the same procedures of virus inoculation, ART, and ATI. They were intravenously infected with 2.2X10^5^ IU SIVmac239M. Beginning at 12 wpi, the animals received a combination of ART consisting of tenofovir disoproxil fumarate (TDF), 5.1 mg/kg; emtricitabine (FTC), 40 mg/kg; and dolutegravir (DTG) 2.5 mg/kg once daily by subcutaneous injection. TDF and FTC were generously provided by Gilead Sciences, Inc. (Foster City, CA), and DTG was generously provided by ViiV Healthcare Limited (Research Triangle, NC). ART continued until 54 wpi when the first ATI occurred from 54 wpi until 57 wpi. This period is referred to as ATI-1. The treatment resumed at 58 wpi and continued until one week before each animal’s necropsy. This 1 week of ATI before necropsy is referred to as ATI-2.

### Sample collection and animal euthanasia

Longitudinal blood samples were obtained and euthanasia was performed following the Tulane IACUC standards of operation and the AVMA Guidelines on the euthanasia of animals. Animals were anesthetized using telazol and buprenorphine, followed by a lethal intravenous injection of sodium pentobarbital. Multiple tissues including MesLNs, IngLNs, spleen, colon, ileum, lung, liver, brain, and blood were collected at necropsy. Fresh samples were stored in snap-frozen or RNAlater during necropsies, or collected for lymphocyte isolation from secondary lymphoid tissues.

### Quantification of vRNA in plasma

Longitudinal blood samples were collected in EDTA-treated tubes. Plasma was separated from whole blood by centrifugation and stored at -80 °C until use. vRNA was extracted from plasma by using the Macherey-Nagel NucleoSpin Virus kit (Catalog No. 740956.50). The pVL was quantified by Q-RT-PCR assay with the primers and probes conserved for the *gag* region of SIVmac239 as described previously (*11*). The primer and probe information is provided in **Table S4**. All samples were measured by Q-RT-PCR with a limit of detection of 81 copies/mL. In addition, at 28, 32, 36, and 40 wpi, and necropsy, samples underwent a more sensitive Q-RT-PCR for evaluation with a limit of detection of 22 copies per mL.

### Quantification of cell-associated SIV DNA and RNA from blood and tissues

RNA and DNA from PBMCs and different tissues were processed as previously described (*11*). Briefly, PBMCs were isolated from blood samples by using Lymphocyte Separation Media (Corning) together with Ammonium-Chloride-Potassium Lysing Buffer (Gibco) for 15min. Frozen PBMCs, lymphocytes from secondary lymphoid tissues, and frozen tissues preserved in RNAlater were used for genomic DNA extraction and RNA extraction. After tissue homogenization using TissueRuptor II (Qiagen) for 30 seconds at medium speed in 30mg of each tissue sample, DNA and RNA were extracted from the dissociated and isolated cells using the AllPrep DNA/RNA kit (Qiagen) following the manufacturer’s instructions. The TaqMan™ Universal PCR Master Mix (Applied Biosystems™) and TaqMan™ RNA-to-CT™ 1-Step Kit (ThermoFisher) were used in the Q-RT-PCR reaction for DNA and RNA samples, respectively. The levels of the cell-associated SIV DNA and CA SIV RNA loads were determined using the methods described earlier using an input of ∼600ng. The number of SIV copies was normalized to diploid genome cell equivalents co-determined by Q-RT-PCR using the TaqMan™ RNase P Detection Reagents Kit (Applied Biosystems), as described previously (*11, 45*). The tissue Q-RT-PCR DNA and RNA assay sensitivity was 10 copies per million cells.

### Deep sequencing of barcoded viral clonotypes

Diagnostic nested PCR was performed on all the samples and positive samples were subjected to deep sequencing as previously described (*7*). A total of 30 mg was processed for DNA/RNA extraction and the same input DNA/RNA was used from each tissue. PCR prior to sequencing was performed with virus-specific primers combined with either the F5 or F7 Illumina adaptors containing unique 8-nucleotide index sequences for multiplexing as previously described (*9*). Sequences were demultiplexed into individual samples based on exact matches to the Illumina P5 index. After index splitting, sequences were aligned to the first 28 bases of the *vpr* gene allowing for 2 nucleotide mismatches. The 34 bases directly upstream from the start codon for *vpr* were extracted, corresponding to the barcode. Single genome amplification followed by Sanger sequencing was used in low-template samples (*7*).

### Fluorescence-activated single-cell sorting and intact proviral DNA assay

CD4+ helper T cells (CD3+, CD4+, CD8-) and myeloid cells (CD3-, CD11b+) were sorted from isolated lymphocytes of spleen and MesLN using the FACSAria III Cell Sorter (BD) together with the following fluorescently-conjugated monoclonal antibody clones from BD: CD8 Sk1 Cy5.5, CD11b ICRF44 PE, CD3 SP34-2 APC, CD4 L200 BV605. Briefly, cells were thawed and kept overnight in a 15 mL conical tube with complete media. Next, the cells were washed twice with PBS containing 0.5% bovine serum albumin and resuspended in FACS buffer. After treating the cells for 15 minutes (min) with Fc Receptor Binding Inhibitor Polyclonal Antibody (ThermoFisher), they were stained for 30 min at 4°C in the dark and washed twice again. An example of the sorting panels used is shown in **Fig. S11**. About 1 million CD4+ T cells and 300,000 myeloid cells were recovered by sorting.

The genomic DNA from sorted cells was extracted as explained above. To perform IPDA, we followed previously described protocols (*46–49*). To detect intact SIV proviral genomes, we used the primer set described by Mancuso et al. (*49*), including the hypermutation probe that did not contain any fluorophore to diminish the IPDA recognition of incompetent virus due to point mutations. We quantified the housekeeping gene *RPP30* using the primers described by Candena et al. (*4*) to quantify the DSI and to normalize the results to SIV copies/10^6^ cells. These numbers were calculated using the methods described by Levy et al (*48*). Approximately 800 ng of gDNA was combined with 600nM of each primer, 200 nM of each probe, and ddPCR supermix for Probes (No dUTP) (Bio-Rad) in the final reaction. PCR was performed following the manufacturer’s instructions.

Approximately 5ng was used to measure the *RRP30* gene in each sample. DSI was used to account for the level of genomic fragmentation of SIV genomes that were not related to defective viral particles. This was accomplished by determining the fragmentation of the *RPP30* gene between the binding sites of the two probes, being the distance between these two probes similar to the distance between the SIV probes *pol* and *env*. After the DSI was calculated, the number of viral copies was normalized to copy numbers per 10^6^ cells. All the primers and probes used are listed in **Table S4.**

SIV biological negative samples and NTC samples were used to set the thresholds for SIV and *RPP30*, respectively. All the samples were run in duplicate and the collective result is shown. The samples reported had >10,000 droplet counts and the SIV-positive samples had a DSI below 0.38 (**Fig. S12**). The data was analyzed using the QuantSoft software Vr. 1.7.4.0917 (Bio-Rad).

### RNAscope and Immuno-fluorescence

RNAscope was conducted using RNAscope 2.5 HD assay kit (Advanced Cell Diagnostics) as reported earlier (*50*) with some modifications. Briefly, the deparaffinized tissue sections were treated with hydrogen peroxide for 10 min at room temperature (RT), then with 0.3 M hydrochloric acid for 20 min at RT, heated with 1X antigen retrieval reagent at 95°C for 20 min using HybeZ hybridization system, rinsed, and treated with protease plus for 15 min at 40°C. The tissues were then incubated with an SIV probe targeting *gag* and *pol* sense strands of SIVmac239 (Advanced Cell Diagnostics) for 2 h at 40°C and washed with 1x SSC buffer (Across Organics) before applying the amplifiers. A series of incubations with the amplifiers 1 to 6 were performed following the manufacturer’s protocol specifications. The tissues were covered in signal detection reagent (diluted 1:60) and incubated at RT. The reaction was stopped with milli-Q water when a visible signal developed and counterstained with 50% hematoxylin (StatLab) for 8 min at RT, rinsed and treated with 0.02% Ammonium hydroxide solution (Sigma) for 30 seconds. Slides were scanned using Zeiss Axio Scan Z.1 (Zeiss) and analyzed using HALO multiplex IHC module (version 3.4 Indica labs).

To characterize the SIV-infected cells, we performed an immunofluorescence assay in combination with RNAscope. The tissue section, after the amplifier treatment mentioned above, was blocked with 10% normal goat serum (Abcam) for 40 min at RT. The tissue was then incubated overnight at 4°C with anti-CD4 (Dako) in a 1:10 dilution, and anti-CD68/CD206 in a 1:20/1:100 dilution (Dako/Sigma). After washing, the tissue section was stained with Alexa Fluor 488 and Alexa Fluor 647 conjugated secondary antibodies in a 1:1000 dilution (Abcam). The tissue was finally counterstained with nuclear stain DAPI in a 1:5000 dilution (EMD Millipore) for 5 min at RT. Images were taken at 63X using the Stellaris Confocal Microscope (Leica).

### Combination of CODEX and RNAscope *in situ* hybridization

Formalin-fixed paraffin-embedded (FFPE) MesLN tissues collected from the necropsied animals LC31, LB66, DJ01, and EM77 were studied using the combination of CODEX and RNAscope *In situ* hybridization (Comb-CODEX-RNAscope) by following our reported method (*51*). Briefly, a 6-μm tissue section on a coverslip was deparaffinized, hydrated, and pretreated with 3% hydrogen peroxide for 10 min at RT, boiling citrate buffer (pH 6, Sigma 21545) for 15 min, and RNAscope® Protease Plus (ACD) at 40°C for 20 min. The tissue section was stained with a cocktail of DNA-barcoded antibodies purchased from Akoya company (CD4/EPR6855, CD20/L26, CD68/KP1, Ki67/B56, CD21/EP3093, CD31/EP3095, HLA-DR/EPR3692) and home-conjugated by us (CD3/SP162). The tissue section on the coverslip was loaded into the CODEX instrument (Akoyo Biosciences) for multiple-cycle immunostaining and image acquisition. After the completion of the CODEX, the tissue-coverslip was subjected to RNAscope procedure with RNAscope® Probe-SIVmac239 (anti-sense, ACD). The CODEX plus RNAscope data were processed by CODEX processor software. The vRNA distribution in relation to T cell and B cell zones was conducted with Voronoi spatial analysis (min:3 µm max:30 µm) using CODEX Multiplex Analysis program (Akoya).

### Single-cell RNA-seq and analysis

Live cells from isolated lymphocytes from MesLN were sorted in FACSAria III Cell Sorter (BD) using LIVE/DEAD Fixable Aqua Dead Cell Stain Kit (Invitrogen). Ten thousand cells were further processed for single-cell sequencing RNA preparation following the instructions of the Chromium Single Cell 3’ Reagent Kits User Guide (v3.1 Chemistry Dual Index, 10x Genomics) to prepare a gene expression library. Libraries were sequenced using Novaseq PE150 (Novogene) with a sequencing depth of 50,000 read pairs per cell. Fastq files were processed with Cellranger (version 7.0.0), using the rhesus macaque reference genome Mmul_10.106 (Ensembl) combined with the SIVmac239 genome (using NBCI reference sequence M33262) divided into 5 sections (SIV_A to SIV-E)^1,2^. The single-cell RNA sequencing dataset was processed, examined, and visualized using Cellenics® community instance (https://scp.biomage.net/) hosted by Biomage (https://biomage.net/).

The following processing settings were used in Cellenics: droplets with a False Discovery Rate (FDR) value < 0.01 based on the emptyDrops method (*52*) were identified as non-empty droplets and retained. Outliers in the distribution of the number of genes vs the number of UMIs were removed by fitting a linear regression model (*p*-values between 7.91E-05 – 2.70E-04). Cells with a high probability of being doublets were filtered out using the scDblFinder method (threshold range: 0.41 – 0.51). Data was log-normalized, and the top 2000 highly variable genes were selected based on the variance stabilizing transformation (VST) method. Reciprocal principal-component analysis (RPCA) was performed, and the top 30 principal components (PCs) were used for data integration using Seurat v4. Finally, to visualize results, a Uniform Manifold Approximation and Projection (UMAP) embedding was calculated. Clusters were identified using the Louvain method, setting an initial resolution of 0.1 to identify main cell types corresponding to large clusters (**Supplementary Table 5**), and a final resolution of 2.0 to further annotate cell subsets within the previously identified cell types (**Supplementary Table 6**). None of the unidentified clusters show any DEG between groups.

### Statistical analysis

Unpaired T-test was used to compare SIV RNA copies per million cells through RNAscope technique using the HALO software in spleen and lymph nodes between the groups. Paired T-test was used to compare the intact provirus by IPDA between spleen and MesLN. The Spearman correlation was used to assess the correlation studies. GraphPad Prism 9.0.1 statistical software (GraphPad Software, Inc., San Diego, CA, USA) was used to analyze data, and statistical results were set to two-sided at *p* < 0.05 as significant.

## Supplementary Materials

Fig. S1 to S12

Table S1 to S6

References *(53-86)*.

## Acknowledgments

The authors would like to thank the animal care staff for their technical assistance. We also thank R. Geleziunas and Gilead Sciences, Inc., as well as C. Parry and ViiV Healthcare for generously providing the antiretroviral drugs for antiretroviral therapy.

## Author contributions

Conceptualization: BL

Methodology: AS-L, NB, SM, FW, GT, YC, XA, VS, CMF, JDL, BFK

Software analysis: AS-L, YL

Validation: AS-L, BL

Formal analysis: AS-L, N B, S M, YC

Data curation: AS-L

Writing-original draft preparation: AS-L

Critical review and discussion: B L, QL, JD L, BFK

Writing-review and revision: AS-L, BFK, BL

Supervision: BL

Project administration: BL

Funding acquisition: BL

## Data and materials availability

The read counts, and Seurat objects from the single-cell RNA sequencing analysis are available in BioStudies under the accession number S-BSST975.

