## Supplementary material for "Lymphoid tissues contribute to viral clonotypes present in plasma at early post-ATI in SIV-infected rhesus macaques": Figures S1-S12 Tables S1-S6

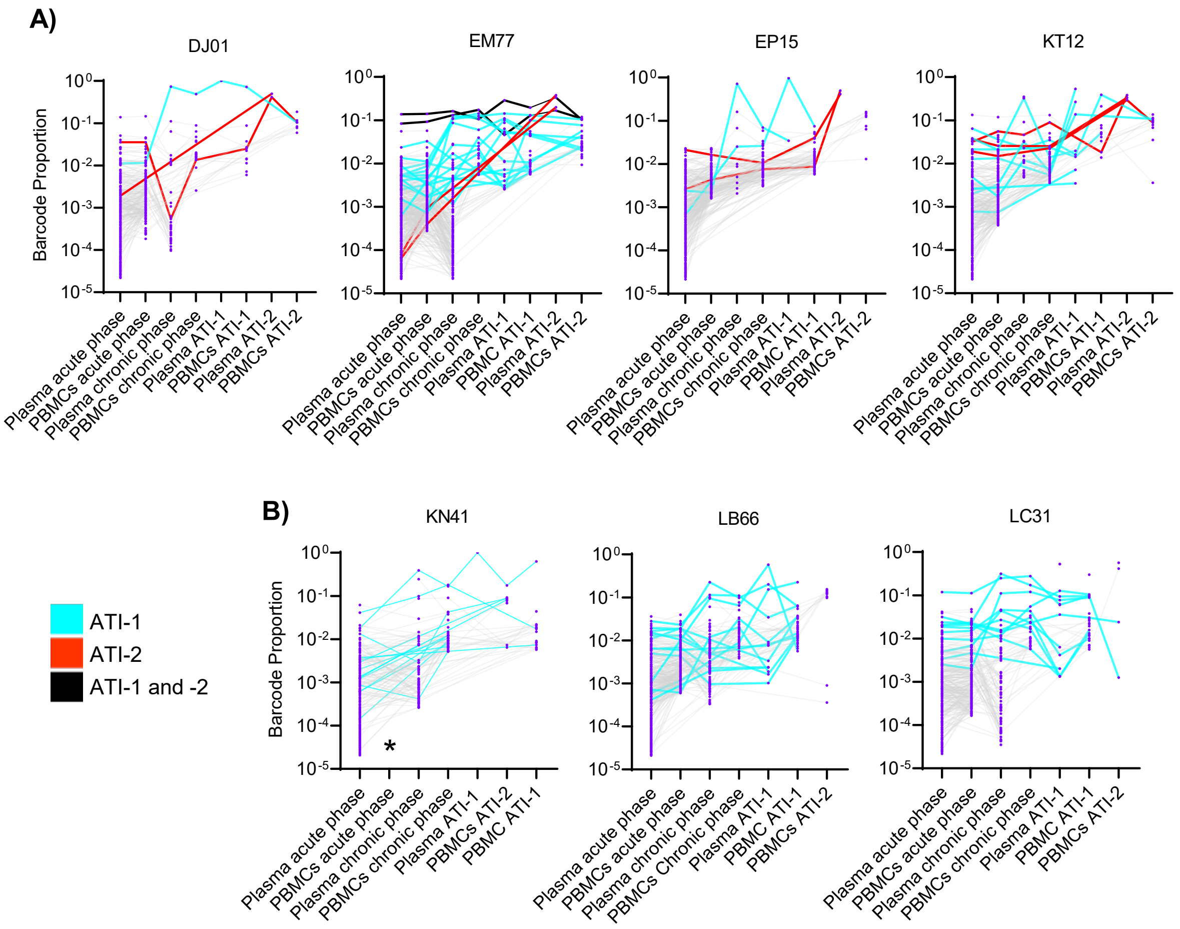


**Figure S1: Detection of SIVmac239M barcodes in CA-DNA from PBMCs.** The barcodes detected in plasma during ATI-1 and ATI-2 were included as a reference. Barcodes associated only with ATI-1 are shown with cyan lines. Barcodes associated only with ATI-2 are shown with red lines. Barcodes associated with ATI-1 and 2 are shown with black lines. **A)** Group 1. **B)** Group 2. *: unavailable.

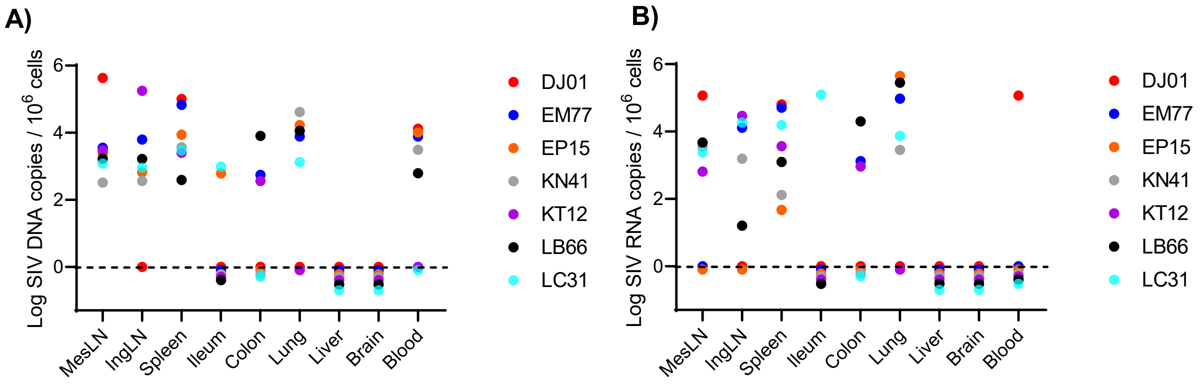


**Figure S2: Levels of cell-associated viral DNA and RNA from the blood and tissues at ATI-2.** Viral DNA and RNA were undetectable in samples on or below dashed line and thus are listed for the visualization of all the tested individual samples. **A**) Cell-associated viral DNA, and **B**) Cell-associated viral RNA was quantified and normalized to copies/million cells from each specific tissue.

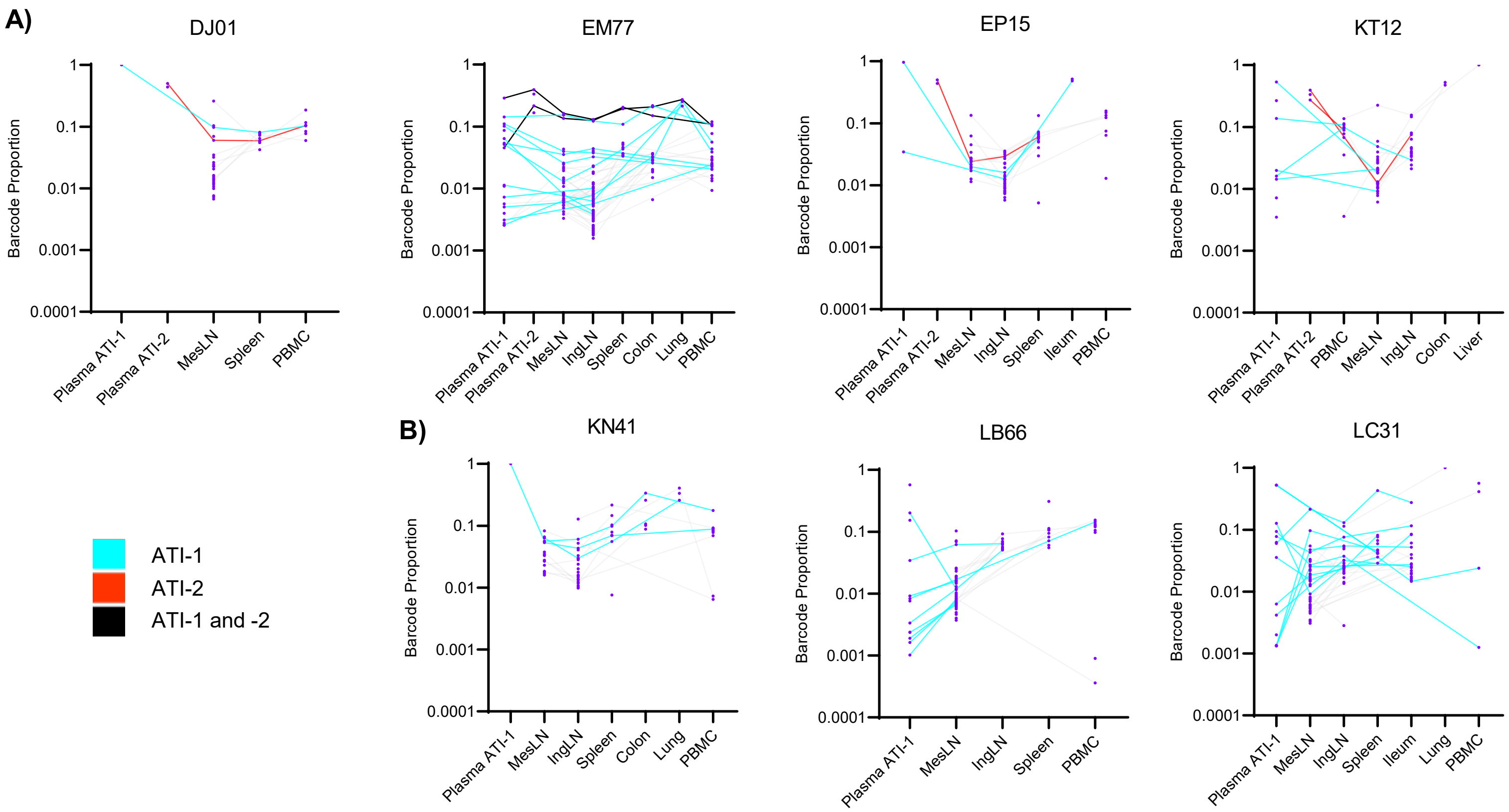


**Figure S3: SIVmac239M barcode detection in cell-associated DNA (CA-DNA) of multiple tissues corresponding with plasma barcodes detected at ATI-1 and ATI-2.** The plasma RNA from ATI-1 and ATI-2 were included as reference of viral clonotypes related to rebound. Barcodes associated only with ATI-1 are shown with a cyan line. Barcodes associated only with ATI-2 are shown with a red line. Barcodes associated with ATI-1 and -2 are shown with a black line. **A)** Group 1. **B)** Group 2.

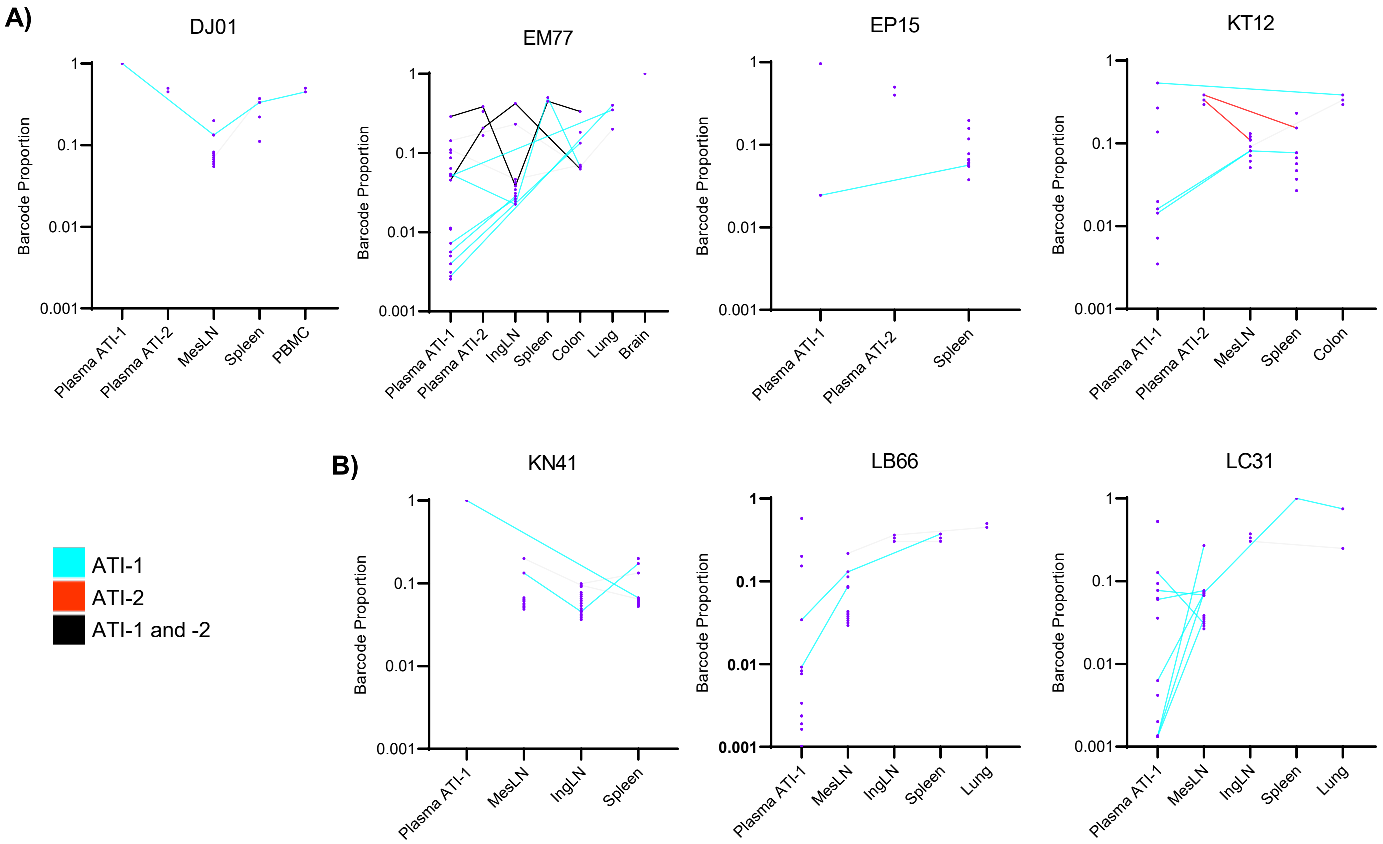


**Figure S4: SIVmac239M barcode detection in cell-associated RNA (CA-RNA) of multiple tissues corresponding with plasma barcodes detected at ATI-1 and ATI-2.** The plasma RNA from ATI-1 and ATI-2 were included as reference of viral clonotype related to rebound. Barcodes associated with ATI-1 alone are shown with a cyan line. Barcodes associated with ATI-2 alone are shown with a red line. Barcodes associated with ATI-1 and -2 are shown with a black line. **A)** Group 1. **B)** Group 2.

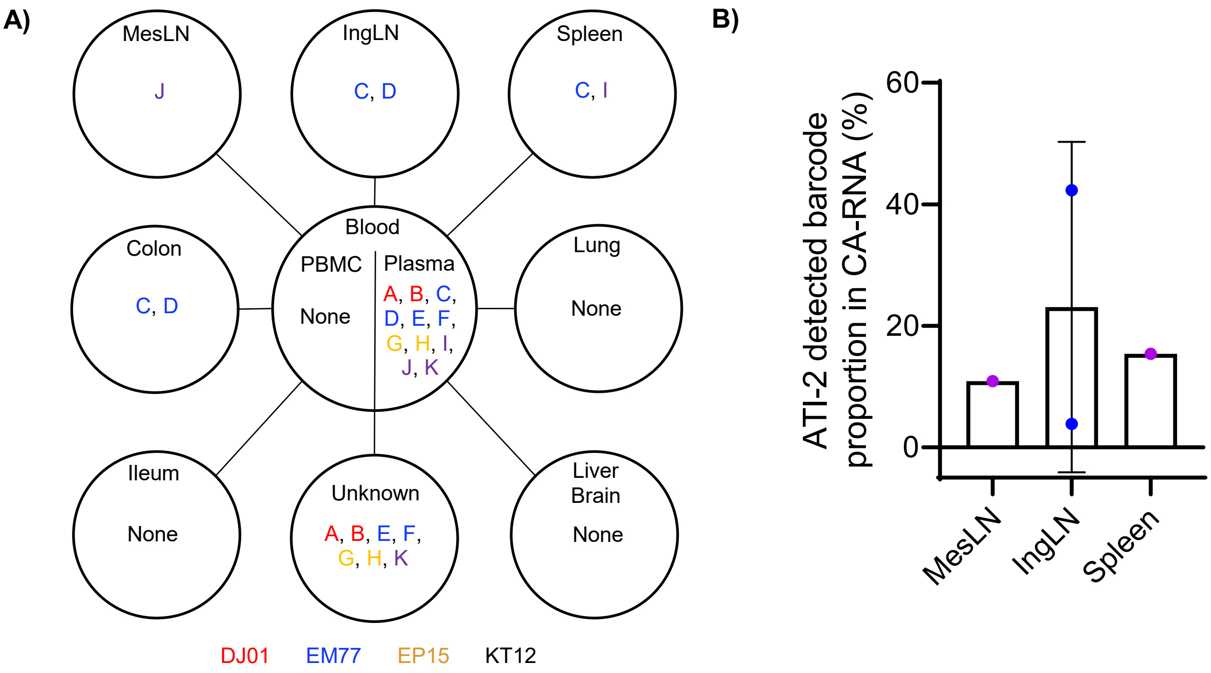


**Figure S5: SIVmac239M barcode detection in CA-RNA from 9 tissues corresponding with plasma barcodes detected at ATI-1 and ATI-2. A)** Barcodes detected from tissue CA-RNA and plasma during ATI-2. They were represented by the letters A to K. Barcodes recovered from each individual are shown as follows: DJ01 (red, A-B), EM77 (blue, C-F), EP15 (gold, G-H) and KT12 (purple, I-K). **B)** Tissue distribution of the total proportion of the ATI-2 detected plasma viral clonotypes.

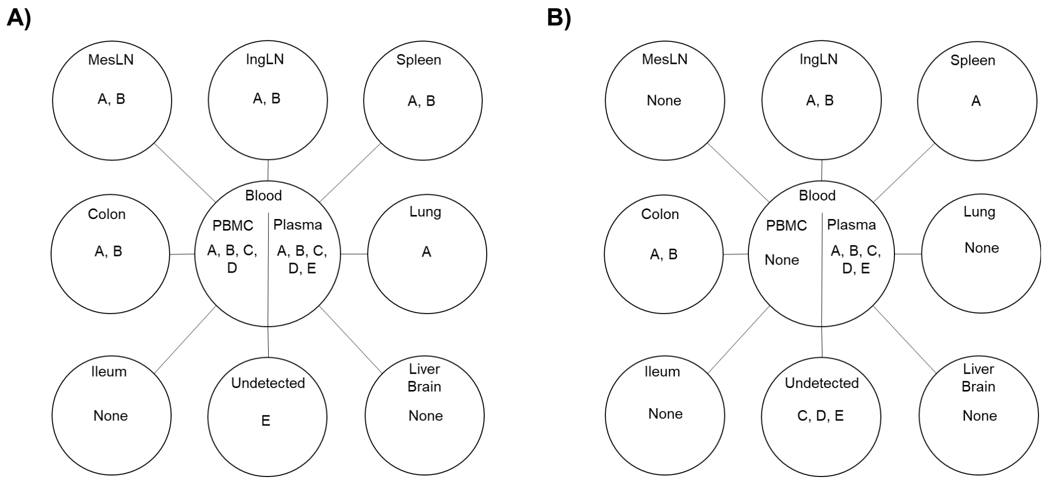


**Figure S6: Barcodes detected in animal KN41 from group 2. A).** Barcodes detected in tissue CA-DNA and in plasma. **B)** Barcodes detected in tissue CA-RNA and in plasma.

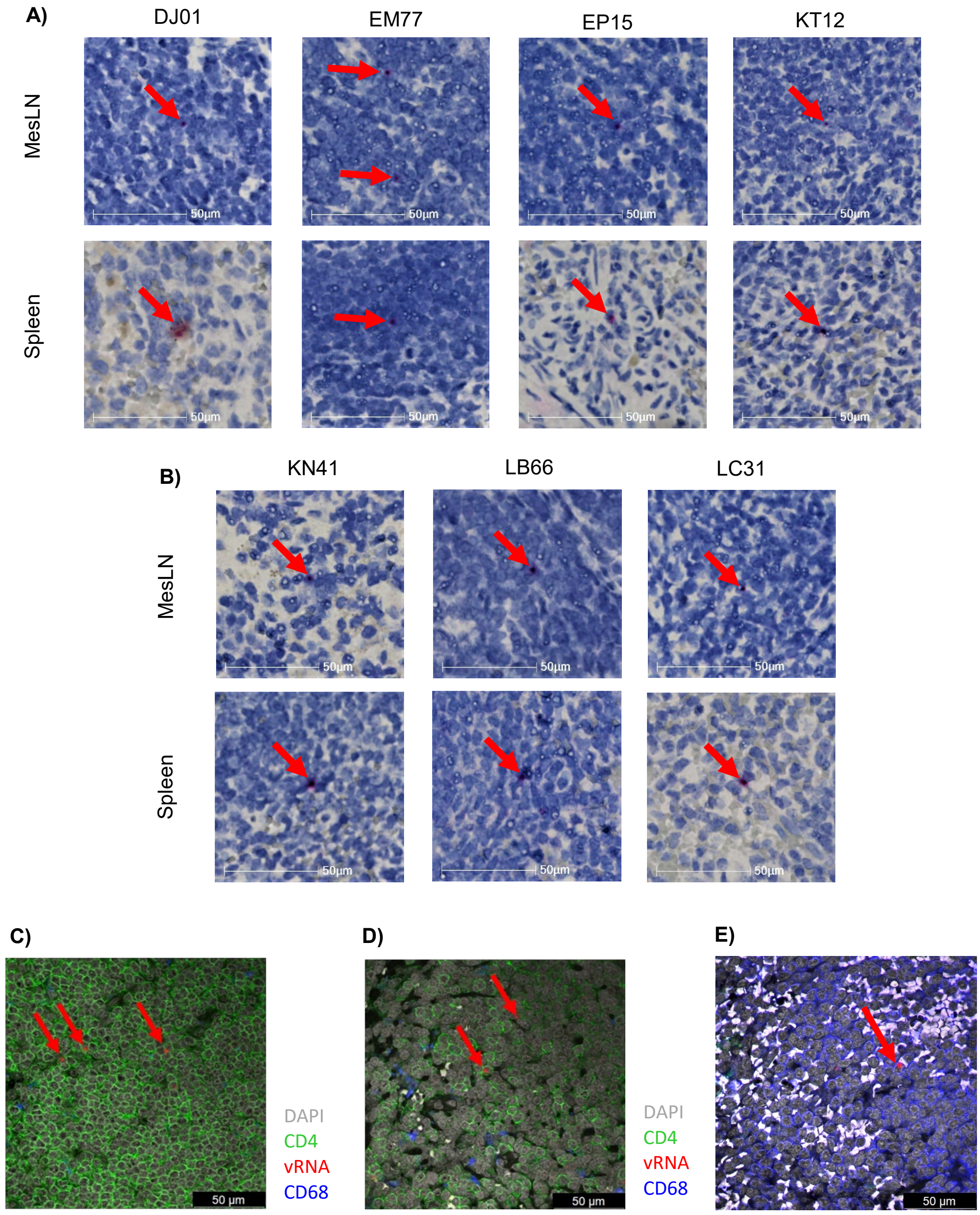


**Figure S7: SIV vRNA positive cells in MesLN and spleen at ATI-2, and colocalization with CD4+ helper T cells and macrophages. A)** group 1. **B)** group 2**. C)** EM77 MesLN at 63X and **D)** EM77 spleen at 63X**. E)** EM77 spleen, red pulp region.

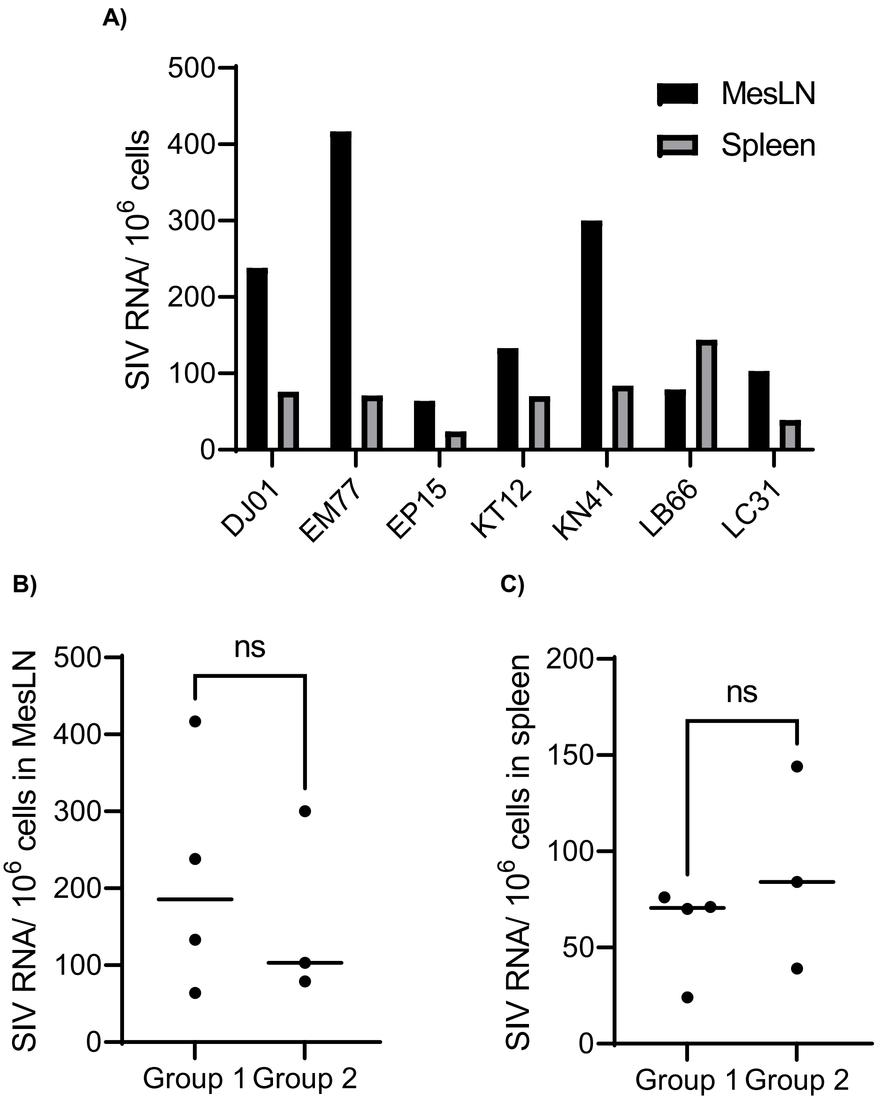


**Figure S8: Levels of SIV RNA positive cells in MesLN and spleen at ATI-2. A**) Individual animal SIV RNA positive cells per million cells in MesLN and Spleen. **B**) Comparison in MesLN between group 1 and group 2. **C**) Comparison in spleen between group 1 and group 2.

A)

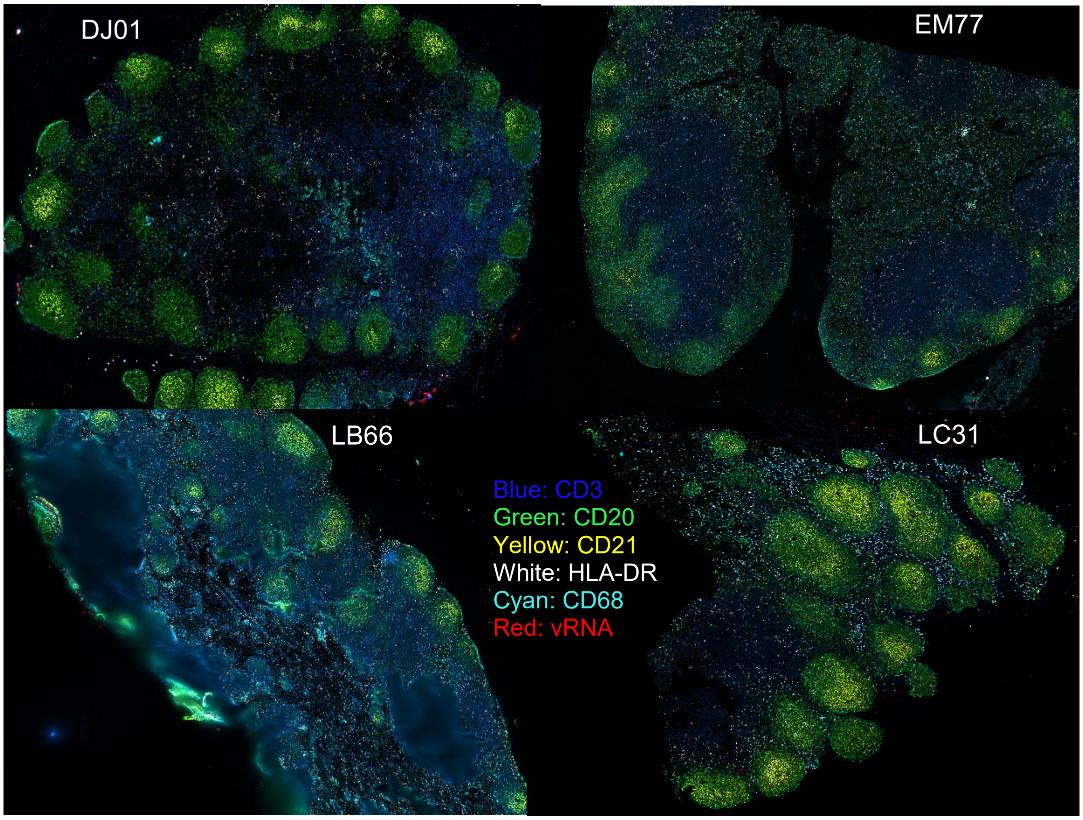


B)

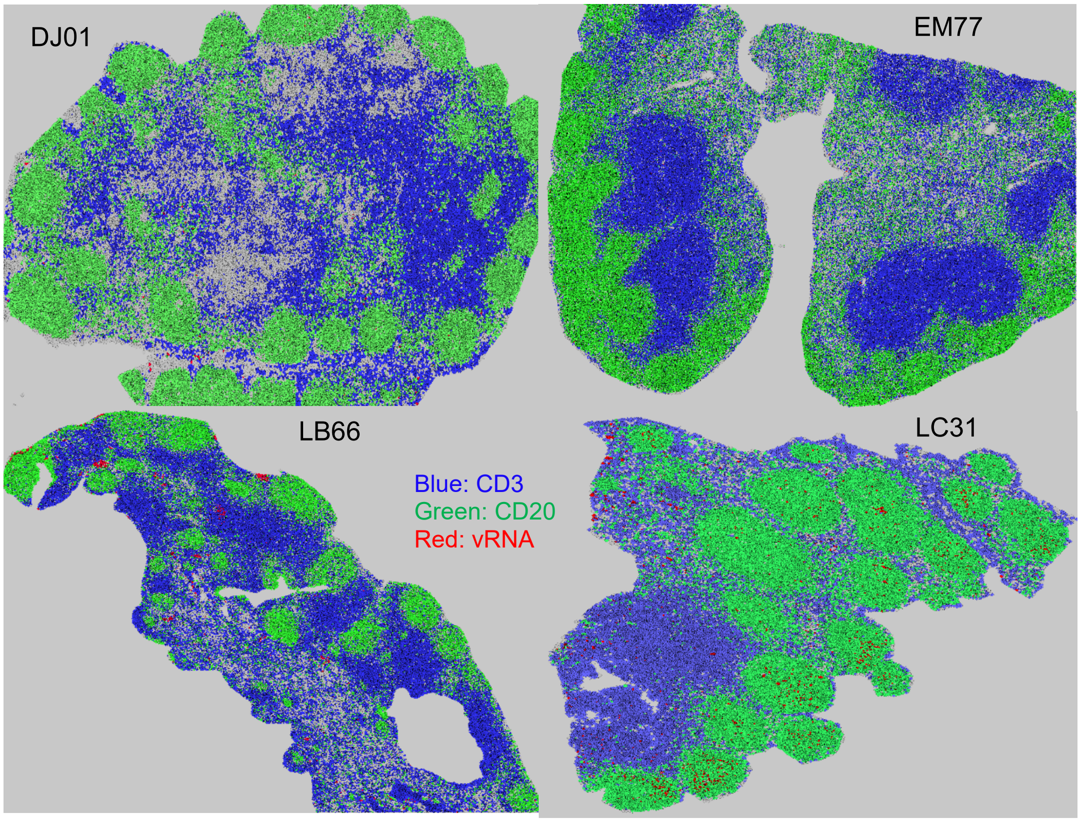


**Figure S9. Com-CODEX-RNAscope from MesLNs at ATI-2.** Images of concurrent detection of multiplexed immunological protein markers and SIV vRNA in MesLN tissues. **A**) A merged image of vRNA, CD68, CD3, CD20, CD21, CD31, HLA-DR from each animal. **B**) A Voronoi plot of the previous figure shows vRNA (in red) distribution in the B cell zones (marked by CD20 in green) and T cell zones (Marked by CD3 in blue).

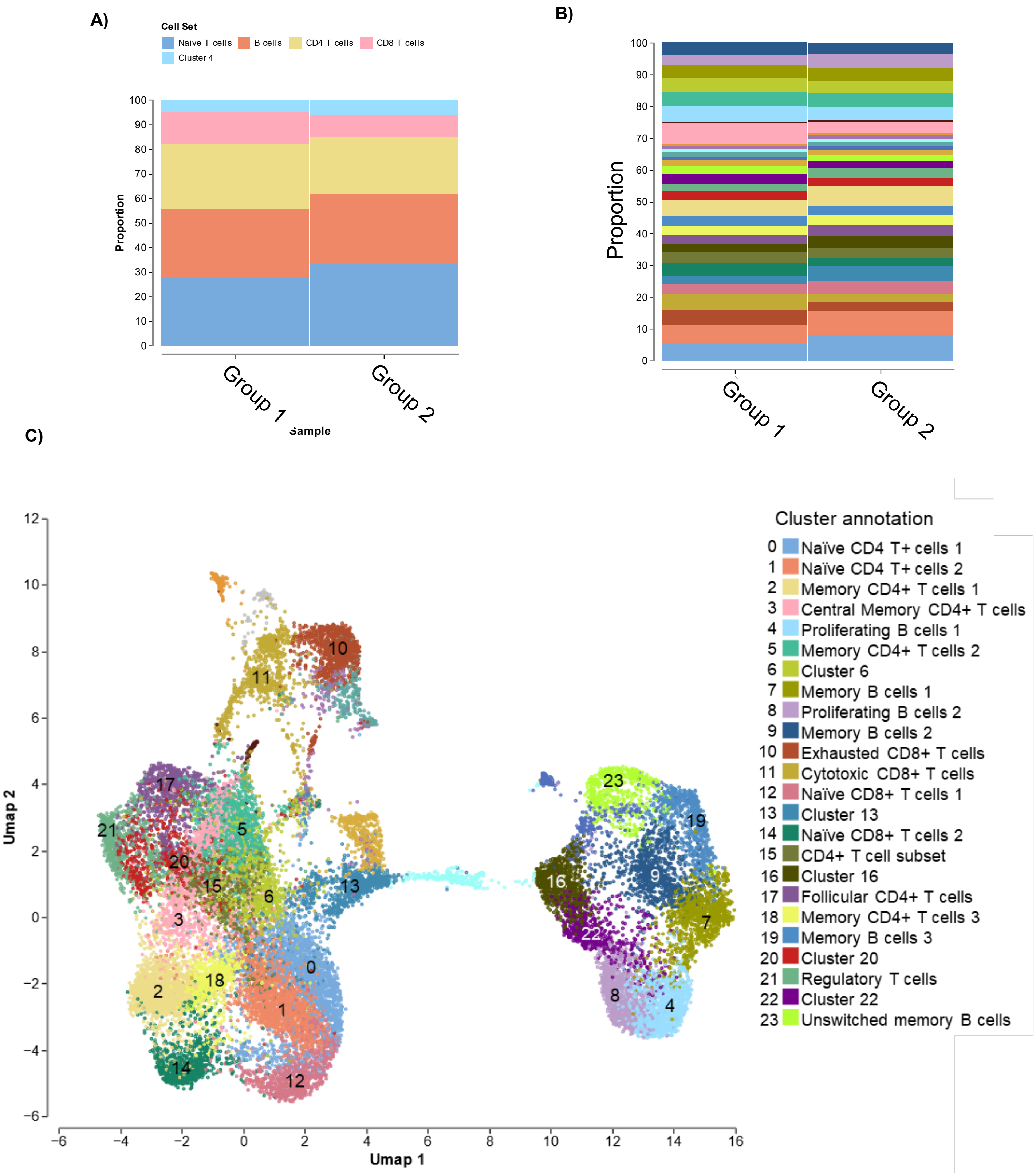


**Figure S10. Additional single-cell RNA sequencing analysis**. **A)** Proportion of louvin clusters at 0.1 resolution. **B)** Proportion of louvin clusters at 2.0 resolution. **C)** Louvin clusters at 2.0 resolution.

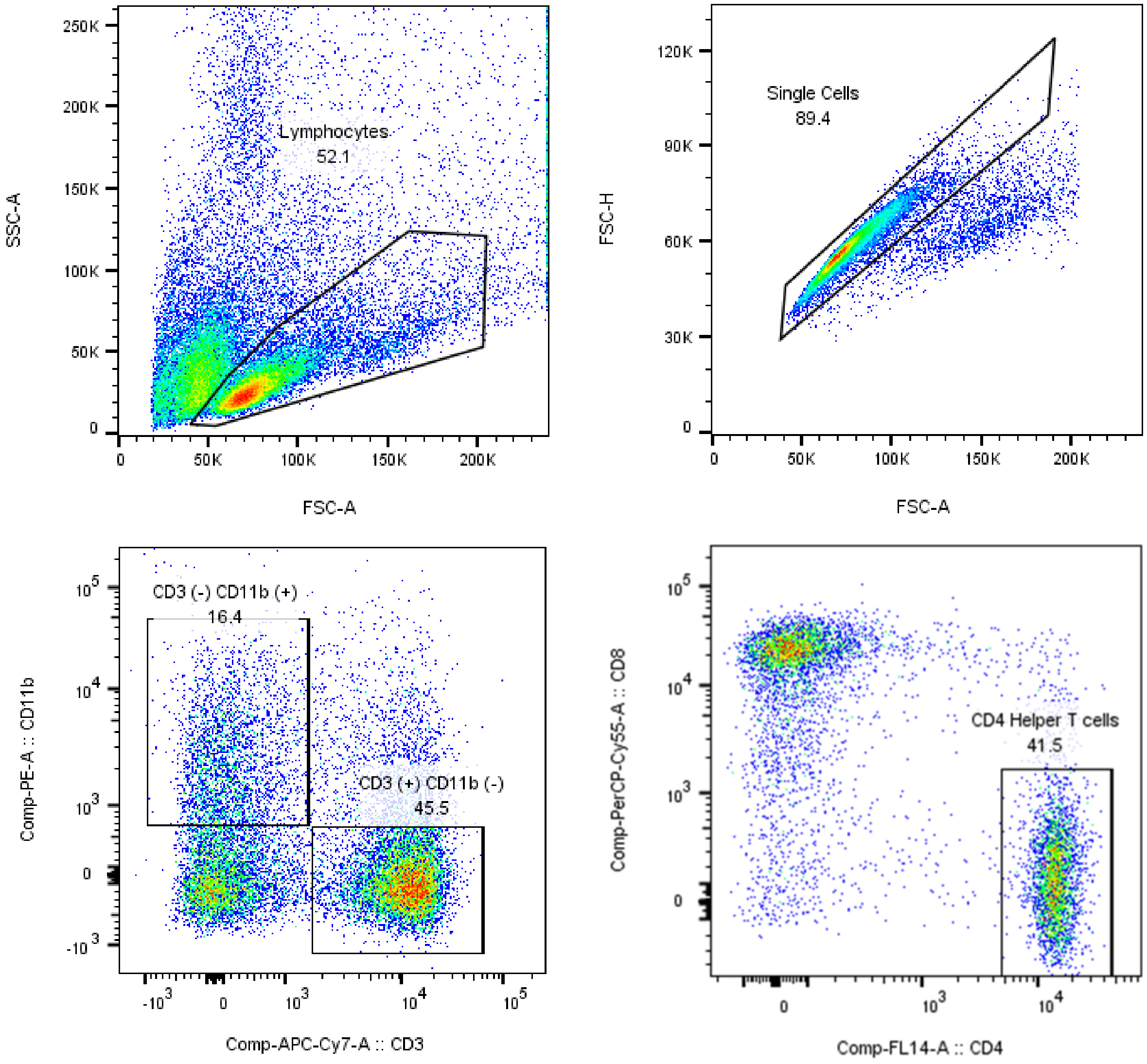


**Figure S11: CD4+ T cells and CD11b+ macrophages sorted from MesLN and spleen.** Cells were gated by forward (FSC) vs side scatter (SSC), then as singlets, then by CD3- CD11b+ for macrophages or CD3+ for T cells and then by CD4+ cells.

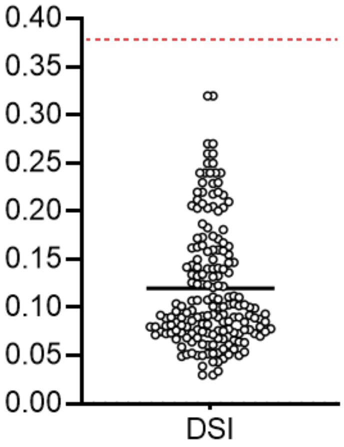


**Figure S12: DNA Shearing Index (DSI) of longitudinal intact proviral sequences in PBMCs and tissues.** RPP30 was measured in each sample to determine the DSI and to normalize the result to intact provirus per $10^6$ PBMCs or $10^6$ CD4+ T cells. The red dashed line indicates the acceptable DSI threshold used.

### Supplementary Table 1: Cluster proportions of scRNAseq analysis

| Louvain Clusters | PBDA2 proportion | Non-PBDA2 proportion |
|---|---|---|
| **Naïve T cells** | 27.563 | 33.257 |
| **B cells** | 27.901 | 28.499 |
| **CD4+ T cells** | 26.619 | 23.216 |
| **CD8+ T cells** | 13.064 | 8.838 |
| **Cluster 4** | 4.854 | 6.189 |

**Note:** Louvin cluster proportions with 0.1 Resolution.

### Supplementary table 2: Highlighted DEGs from scRNAseq

| DEG | Associated relation | Reference |
|---|---|---|
| **COX7C** | Mitochondria dysfunction | 53 |
| **IMMP2L** | Mitochondria dysfunction | 54 |
| **MICOS13** | Mitochondria dysfunction | 55 |
| **CERS6** | Mitochondria dysfunction | 56 |
| **COX1** | Mitochondria dysfunction | 53 |
| **PRDX6** | Oxidative stress | 57 |
| **OXR1** | Oxidative stress | 58 |
| **CERKL** | Oxidative stress | 59 |
| **NXN** | Oxidative stress | 60 |
| **H2AC20** | Nucleosome and histone modifications | 61 |
| **TOX** | Nucleosome and histone modifications/ Immune response | 62, 63 |
| **TET3** | Nucleosome and histone modifications | 64 |
| **KAT6A** | Nucleosome and histone modifications | 65 |
| **H2AC12** | Nucleosome and histone modifications | 61 |
| **FYN** | Nucleosome and histone modifications / Immune response | 66, 67 |
| **PLAC8** | Immune response | 68 |
| **SVIP** | Immune response | 69 |
| **GPR65** | Immune response | 70 |
| **CAMK1D** | Immune response | 71 |
| **RNF130** | Immune response | 72 |
| **ISG15** | Immune response | 73 |
| **TNFSF13B** | Immune response | 74 |
| **FBXO27** | Immune response | 75 |
| **CD1B** | Immune response | 76 |
| **ANXA1** | Immune response | 77 |
| **RGS6** | Immune response | 78 |
| **LGALS3** | Immune response | 79 |
| **CLEC2D** | Immune response | 80 |
| **THEMIS** | Immune response | 81 |
| **KLRB1** | Immune response | 82 |
| **GZMB** | Immune response | 83 |
| **S100A4** | Immune response | 84 |
| **PRF1** | Immune response | 85 |
| **NKG7** | Immune response | 86 |

### Supplementary Table 3: Total list of DEGs per cell subset

| Cluster in 0.1 resolution | Cluster in 2.0 resolution | Cell subset | DEG | LogFC | adj p-value |
|---|---|---|---|---|---|
| **Naïve T cells** | 0 | Naïve CD4+ T cells 1 | CALY | 1.9327 | 0.0358 |
| **Naïve T cells** | 0 | Naïve CD4+ T cells 1 | PLAC8 | 1.648 | 0.0092 |
| **Naïve T cells** | 0 | Naïve CD4+ T cells 1 | COX7C | 1.5665 | 0.0092 |
| **Naïve T cells** | 0 | Naïve CD4+ T cells 1 | H2AC20 | 1.3784 | 0.0286 |
| **Naïve T cells** | 0 | Naïve CD4+ T cells 1 | KCNQ5 | -1.3056 | 0.048 |
| **Naïve T cells** | 1 | Naïve CD4+ T cells 2 | PRELID2 | 3.8098 | 0.0491 |
| **Naïve T cells** | 1 | Naïve CD4+ T cells 2 | GIPC2 | 2.3382 | 0.0071 |
| **Naïve T cells** | 1 | Naïve CD4+ T cells 2 | COX7C | 1.8679 | 0.0223 |
| **Naïve T cells** | 1 | Naïve CD4+ T cells 2 | ITGA2 | 1.8346 | 0.016 |
| **Naïve T cells** | 1 | Naïve CD4+ T cells 2 | CALY | 1.6477 | 0.0288 |
| **Naïve T cells** | 1 | Naïve CD4+ T cells 2 | SVIP | 1.4433 | 0.0326 |
| **Naïve T cells** | 1 | Naïve CD4+ T cells 2 | PLAC8 | 1.3262 | 0.0071 |
| **Naïve T cells** | 1 | Naïve CD4+ T cells 2 | DIP2C | 1.2972 | 0.0071 |
| **Naïve T cells** | 1 | Naïve CD4+ T cells 2 | CYB5D2 | 1.1982 | 0.0342 |
| **Naïve T cells** | 1 | Naïve CD4+ T cells 2 | SUCLG2 | 1.0972 | 0.016 |
| **Naïve T cells** | 1 | Naïve CD4+ T cells 2 | GPR65 | 1.0187 | 0.0407 |
| **Naïve T cells** | 1 | Naïve CD4+ T cells 2 | PRDX6 | 1.0042 | 0.0435 |
| **Naïve T cells** | 1 | Naïve CD4+ T cells 2 | OXR1 | -0.9627 | 0.0344 |
| **Naïve T cells** | 1 | Naïve CD4+ T cells 2 | CAMK1D | -0.997 | 0.0435 |
| **Naïve T cells** | 1 | Naïve CD4+ T cells 2 | SERGEF | -1.0538 | 0.0444 |
| **Naïve T cells** | 1 | Naïve CD4+ T cells 2 | TOX | -1.0761 | 0.0128 |
| **Naïve T cells** | 1 | Naïve CD4+ T cells 2 | IMMP2L | -1.1024 | 0.0176 |
| **Naïve T cells** | 1 | Naïve CD4+ T cells 2 | TET3 | -1.1377 | 0.0407 |
| **Naïve T cells** | 1 | Naïve CD4+ T cells 2 | TANC2 | -1.352 | 0.0071 |
| **Naïve T cells** | 1 | Naïve CD4+ T cells 2 | RNF130 | -1.4813 | 0.0097 |
| **Naïve T cells** | 1 | Naïve CD4+ T cells 2 | C1H1orf159 | -1.6592 | 0.0071 |
| **Naïve T cells** | 1 | Naïve CD4+ T cells 2 | KCNQ5 | -1.7285 | 0.0002 |
| **Naïve T cells** | 1 | Naïve CD4+ T cells 2 | APOD | -1.8542 | 0.0407 |
| **Naïve T cells** | 1 | Naïve CD4+ T cells 2 | COX1 | -2.4765 | 0.0407 |
| **Naïve T cells** | 1 | Naïve CD4+ T cells 2 | ANKS1B | -3.3368 | 0.0407 |
| **Naïve T cells** | 1 | Naïve CD4+ T cells 2 | PTK2 | -4.8075 | 0.0435 |
| **Naïve T cells** | 2 | Memory CD4+ T cells 1 | ABO | 4.0282 | 0.0078 |
| **Naïve T cells** | 2 | Memory CD4+ T cells 1 | GIPC2 | 3.1082 | 0.0208 |
| **Naïve T cells** | 2 | Memory CD4+ T cells 1 | MYO16 | 3.0796 | 0.0021 |
| **Naïve T cells** | 2 | Memory CD4+ T cells 1 | ISG15 | 2.9197 | 0.0021 |
| **Naïve T cells** | 2 | Memory CD4+ T cells 1 | TNFSF13B | 2.2111 | 0.0045 |
| **Naïve T cells** | 2 | Memory CD4+ T cells 1 | SIPA1L2 | 2.2054 | 0.0317 |
| **Naïve T cells** | 2 | Memory CD4+ T cells 1 | CPQ | 2.1208 | 0.0018 |

| | | | | | |
|---|---|---|---|---|---|
| **Naïve T cells** | 2 | Memory CD4+ T cells 1 | CHMP7 | 2.0705 | 0.003 |
| **Naïve T cells** | 2 | Memory CD4+ T cells 1 | FBXO27 | 1.9383 | 0.0339 |
| **Naïve T cells** | 2 | Memory CD4+ T cells 1 | R3HCC1 | 1.7249 | 0.0468 |
| **Naïve T cells** | 2 | Memory CD4+ T cells 1 | H2AC20 | 1.4887 | 0.0021 |
| **Naïve T cells** | 2 | Memory CD4+ T cells 1 | COX7C | 1.4588 | 0.0009 |
| **Naïve T cells** | 2 | Memory CD4+ T cells 1 | CERKL | 1.4211 | 0.0466 |
| **Naïve T cells** | 2 | Memory CD4+ T cells 1 | PLAC8 | 1.3803 | 0.0014 |
| **Naïve T cells** | 2 | Memory CD4+ T cells 1 | MICOS13 | 1.3543 | 0.0045 |
| **Naïve T cells** | 2 | Memory CD4+ T cells 1 | PRDX6 | 1.3483 | 0.003 |
| **Naïve T cells** | 2 | Memory CD4+ T cells 1 | DIP2C | 1.2964 | 0.0094 |
| **Naïve T cells** | 2 | Memory CD4+ T cells 1 | BEX1 | 1.2103 | 0.0339 |
| **Naïve T cells** | 2 | Memory CD4+ T cells 1 | SUCLG2 | 1.1506 | 0.0031 |
| **Naïve T cells** | 2 | Memory CD4+ T cells 1 | EIF3G | 1.0428 | 0.0234 |
| **Naïve T cells** | 2 | Memory CD4+ T cells 1 | GTF2H5 | 0.9781 | 0.044 |
| **Naïve T cells** | 2 | Memory CD4+ T cells 1 | KAT6A | -0.8487 | 0.0353 |
| **Naïve T cells** | 2 | Memory CD4+ T cells 1 | PHACTR2 | -0.9551 | 0.0418 |
| **Naïve T cells** | 2 | Memory CD4+ T cells 1 | C1H1orf159 | -1.4047 | 0.0103 |
| **Naïve T cells** | 2 | Memory CD4+ T cells 1 | PRKN | -1.4102 | 0.0034 |
| **Naïve T cells** | 2 | Memory CD4+ T cells 1 | KCNQ5 | -1.5541 | 0.0001 |
| **Naïve T cells** | 2 | Memory CD4+ T cells 1 | APOD | -1.7944 | 0.0063 |
| **Naïve T cells** | 2 | Memory CD4+ T cells 1 | CD1B | -2.0254 | 0.0229 |
| **Naïve T cells** | 2 | Memory CD4+ T cells 1 | NXN | -2.4519 | 0.008 |
| **Naïve T cells** | 12 | Naïve CD8+ T cells 1 | PLAC8 | 1.6227 | 0.0371 |
| **Naïve T cells** | 14 | Naïve CD8+ T cells 2 | DNAJB1 | 3.3869 | 0 |
| **Naïve T cells** | 14 | Naïve CD8+ T cells 2 | TNFSF13B | 2.8802 | 0.0045 |
| **Naïve T cells** | 14 | Naïve CD8+ T cells 2 | ANXA1 | 2.4536 | 0.0291 |
| **Naïve T cells** | 14 | Naïve CD8+ T cells 2 | RGS1 | 2.2878 | 0.044 |
| **Naïve T cells** | 14 | Naïve CD8+ T cells 2 | BMPR1B | 1.7949 | 0.0145 |
| **Naïve T cells** | 14 | Naïve CD8+ T cells 2 | COX7C | 1.6545 | 0.0005 |
| **Naïve T cells** | 14 | Naïve CD8+ T cells 2 | PRDX6 | 1.607 | 0.0066 |
| **Naïve T cells** | 14 | Naïve CD8+ T cells 2 | H2AC20 | 1.5814 | 0.0171 |
| **Naïve T cells** | 14 | Naïve CD8+ T cells 2 | MICOS13 | 1.4177 | 0.029 |
| **Naïve T cells** | 14 | Naïve CD8+ T cells 2 | ZNF483 | 1.0331 | 0.04 |
| **Naïve T cells** | 14 | Naïve CD8+ T cells 2 | ADAMTS6 | -1.0022 | 0.04 |
| **Naïve T cells** | 14 | Naïve CD8+ T cells 2 | TANC2 | -1.4382 | 0.04 |
| **Naïve T cells** | 14 | Naïve CD8+ T cells 2 | KCNQ5 | -1.4661 | 0.0005 |
| **Naïve T cells** | 18 | Memory CD4+ T cells 3 | COX7C | 1.4497 | 0.0247 |
| **Naïve T cells** | 18 | Memory CD4+ T cells 3 | KCNQ5 | -1.409 | 0.0247 |
| **CD4+ T cells** | 3 | Central Memory CD4+ T cells | MYO16 | 2.7975 | 0.0253 |
| **CD4+ T cells** | 3 | Central Memory CD4+ T cells | TNFSF13B | 2.0356 | 0.0367 |
| **CD4+ T cells** | 3 | Central Memory CD4+ T cells | H2AC20 | 1.7849 | 0.0319 |

| | | | | | |
|---|---|---|---|---|---|
| **CD4+ T cells** | 3 | Central Memory CD4+ T cells | COX7C | 1.6667 | 0.0155 |
| **CD4+ T cells** | 3 | Central Memory CD4+ T cells | PDE7B | -1.8617 | 0.0025 |
| **CD4+ T cells** | 3 | Central Memory CD4+ T cells | KCNQ5 | -2.5432 | 0.0006 |
| **CD4+ T cells** | 3 | Central Memory CD4+ T cells | COX1 | -2.5991 | 0.0367 |
| **CD4+ T cells** | 3 | Central Memory CD4+ T cells | RGS6 | -5.4206 | 0.0308 |
| **CD4+ T cells** | 5 | Memory CD4+ T cells 2 | COX7C | 1.8014 | 0.0013 |
| **CD4+ T cells** | 5 | Memory CD4+ T cells 2 | H1-4 | 1.4536 | 0.0409 |
| **CD4+ T cells** | 5 | Memory CD4+ T cells 2 | H2AC20 | 1.2981 | 0.0409 |
| **CD4+ T cells** | 5 | Memory CD4+ T cells 2 | SPAG16 | -2.0233 | 0.0207 |
| **CD4+ T cells** | 5 | Memory CD4+ T cells 2 | RGS6 | -2.9519 | 0.0002 |
| **CD4+ T cells** | 5 | Memory CD4+ T cells 2 | KCNQ5 | -3.8518 | 0.0409 |
| **CD4+ T cells** | 15 | CD4+ T cell subset | CHMP7 | 2.1881 | 0.0495 |
| **CD4+ T cells** | 15 | CD4+ T cell subset | DIP2C | 1.537 | 0.0495 |
| **CD4+ T cells** | 15 | CD4+ T cell subset | COX7C | 1.4733 | 0.0495 |
| **CD4+ T cells** | 15 | CD4+ T cell subset | PDE7B | -1.6699 | 0.0087 |
| **CD4+ T cells** | 17 | Follicular CD4+ T cell | SPINK2 | 4.3688 | 0.0369 |
| **CD4+ T cells** | 17 | Follicular CD4+ T cell | AFDN | 2.5403 | 0.0445 |
| **CD4+ T cells** | 17 | Follicular CD4+ T cell | LGALS3 | 2.073 | 0.0369 |
| **CD4+ T cells** | 17 | Follicular CD4+ T cell | H2AC20 | 1.9474 | 0.0009 |
| **CD4+ T cells** | 17 | Follicular CD4+ T cell | H1-4 | 1.826 | 0.0009 |
| **CD4+ T cells** | 17 | Follicular CD4+ T cell | CHMP7 | 1.8238 | 0.0009 |
| **CD4+ T cells** | 17 | Follicular CD4+ T cell | COX7C | 1.4036 | 0.0034 |
| **CD4+ T cells** | 17 | Follicular CD4+ T cell | CLEC2D | -1.0979 | 0.0314 |
| **CD4+ T cells** | 17 | Follicular CD4+ T cell | IGF1R | -1.245 | 0.0299 |
| **CD4+ T cells** | 17 | Follicular CD4+ T cell | PAM | -1.2912 | 0.0069 |
| **CD4+ T cells** | 17 | Follicular CD4+ T cell | ZDHHC14 | -1.3653 | 0.0399 |
| **CD4+ T cells** | 17 | Follicular CD4+ T cell | MRTFB | -1.3855 | 0.0177 |
| **CD4+ T cells** | 17 | Follicular CD4+ T cell | ARMC2 | -1.3885 | 0.0369 |
| **CD4+ T cells** | 17 | Follicular CD4+ T cell | SCLT1 | -1.4278 | 0.0246 |
| **CD4+ T cells** | 17 | Follicular CD4+ T cell | SNTB1 | -1.9788 | 0.0053 |
| **CD4+ T cells** | 17 | Follicular CD4+ T cell | PARD3 | -2.0835 | 0.0132 |
| **CD4+ T cells** | 17 | Follicular CD4+ T cell | COX1 | -2.1107 | 0.0082 |
| **CD4+ T cells** | 17 | Follicular CD4+ T cell | C1H1orf159 | -2.735 | 0.0177 |
| **CD4+ T cells** | 17 | Follicular CD4+ T cell | SYT14 | -3.8521 | 0.0404 |
| **CD4+ T cells** | 17 | Follicular CD4+ T cell | NRP1 | -4.7091 | 0.0369 |
| **CD4+ T cells** | 21 | Regulatory T cells | S100A4 | 4.008 | 0.031 |
| **CD8+ T cells** | 10 | Exhausted CD8+ T cells | LARGE1 | 3.5743 | 0.0066 |
| **CD8+ T cells** | 10 | Exhausted CD8+ T cells | COX7C | 1.3154 | 0.0222 |
| **CD8+ T cells** | 10 | Exhausted CD8+ T cells | THEMIS | -1.1732 | 0.0476 |
| **CD8+ T cells** | 10 | Exhausted CD8+ T cells | KLRB1 | -1.4905 | 0.0044 |
| **CD8+ T cells** | 10 | Exhausted CD8+ T cells | KCNQ5 | -2.668 | 0.0002 |

| | | | | | |
|---|---|---|---|---|---|
| **CD8+ T cells** | 11 | Cytotoxic CD8+ T cells | GZMB | 3.6264 | 0.0449 |
| **CD8+ T cells** | 11 | Cytotoxic CD8+ T cells | S100A4 | 2.0627 | 0.0014 |
| **CD8+ T cells** | 11 | Cytotoxic CD8+ T cells | PRF1 | 2.0083 | 0.0325 |
| **CD8+ T cells** | 11 | Cytotoxic CD8+ T cells | H2AC20 | 1.7227 | 0.0068 |
| **CD8+ T cells** | 11 | Cytotoxic CD8+ T cells | RYR2 | 1.4336 | 0.0295 |
| **CD8+ T cells** | 11 | Cytotoxic CD8+ T cells | COX7C | 1.3744 | 0.0295 |
| **CD8+ T cells** | 11 | Cytotoxic CD8+ T cells | NKG7 | 1.2576 | 0.0295 |
| **CD8+ T cells** | 11 | Cytotoxic CD8+ T cells | CD52 | 1.2557 | 0.031 |
| **CD8+ T cells** | 11 | Cytotoxic CD8+ T cells | CIB1 | 1.2095 | 0.0456 |
| **CD8+ T cells** | 11 | Cytotoxic CD8+ T cells | CERS6 | -1.191 | 0.0295 |
| **CD8+ T cells** | 11 | Cytotoxic CD8+ T cells | BMP2K | -1.1996 | 0.0449 |
| **CD8+ T cells** | 11 | Cytotoxic CD8+ T cells | PDZRN3 | -1.5528 | 0.0026 |
| **CD8+ T cells** | 11 | Cytotoxic CD8+ T cells | KLRB1 | -1.9605 | 0.0236 |
| **CD8+ T cells** | 11 | Cytotoxic CD8+ T cells | KCNQ5 | -3.5181 | 0.0001 |
| **B cells** | 4 | Proliferating B cells 1 | FYN | -1.8645 | 0.0284 |
| **B cells** | 7 | Memory B cells 1 | H2AC12 | 2.2169 | 0.0261 |
| **B cells** | 7 | Memory B cells 1 | H2AC20 | 1.6008 | 0.0195 |
| **B cells** | 7 | Memory B cells 1 | S100A10 | 1.5016 | 0.0218 |
| **B cells** | 7 | Memory B cells 1 | COX7C | 1.4085 | 0.0287 |
| **B cells** | 7 | Memory B cells 1 | KCNQ5 | -1.3657 | 0.0195 |
| **B cells** | 7 | Memory B cells 1 | CPA6 | -1.4201 | 0.0195 |
| **B cells** | 7 | Memory B cells 1 | CPEB4 | -1.5834 | 0.0261 |
| **B cells** | 7 | Memory B cells 1 | TMEM117 | -1.6428 | 0.0421 |
| **B cells** | 7 | Memory B cells 1 | ITGA1 | -1.7011 | 0.0195 |
| **B cells** | 7 | Memory B cells 1 | COX1 | -2.1552 | 0.0195 |
| **B cells** | 8 | Proliferating B cells 2 | COX7C | 1.4415 | 0.0319 |
| **B cells** | 8 | Proliferating B cells 2 | DEXI | 1.244 | 0.0431 |
| **B cells** | 8 | Proliferating B cells 2 | FYN | -1.2875 | 0.0319 |
| **B cells** | 8 | Proliferating B cells 2 | CPA6 | -1.8544 | 0.0319 |
| **B cells** | 8 | Proliferating B cells 2 | ITGA1 | -5.1451 | 0.0319 |
| **B cells** | 19 | Memory B cells 3 | ITGA1 | -3.1017 | 0.0074 |
| **B cells** | 23 | Unswitched memory B cells (F) | ITGA1 | -2.5467 | 0.0354 |

### Supplementary Table 4: Primers and probes list

| Primer name | Assay | Sequence (5' → 3') |
| --- | --- | --- |
| **Gag Forward** | qRT-PCR Viral Load Assay, CA viral DNA/RNA qPCR Assays | AGGCTGCAGATTGGGACTTG |
| **Gag Reverse** | qRT-PCR Viral Load Assay, CA viral DNA/RNA qPCR Assays | TGATCCTGACGGCTCCCTAA |
| **Gag Prove** | qRT-PCR Viral Load Assay, CA viral DNA/RNA qPCR Assays | FAM-ACCCACAAC-ZEN-CAGCTCCACAACAAGGAC-3IABkFQ |
| **Ψ Forward** | SIV IPDA | ACGACGGAGTGCTCCTATAA |
| **Ψ Reverse** | SIV IPDA | TCACGCCCATCTCCCACTCT |
| **Ψ probe** | SIV IPDA | FAM-TTGTGTTGCACTTACCTGCA-3IABkFQ |
| **RRE Forward** | SIV IPDA | CTACTGGTGGCACCTCAAGA |
| **RRE Reverse** | SIV IPDA | AGCGGTCAGCGTCAACGA |
| **RRE probe** | SIV IPDA | HEX-TGTGCTAGGGTTCTTGGGTT-3IABkFQ |
| **RRE/hyper probe** | SIV IPDA | TGTGCTAAGGTTCTTAGGTT-3IABkFQ |
| **RM RPP30 Forward 1** | SIV IPDA | AGGATGCTCCGGGAGTATGTA |
| **RM RPP30 Reverse 1** | SIV IPDA | CCTGCTTGTCACCTATATAACAT |
| **RM RPP30 Probe 1** | SIV IPDA | FAM-TCAAGCTGGGAGACGGAAGAGTCAGT-3IABkFQ |
| **RM RPP30 Forward 2** | SIV IPDA | ACAGACTCACACAATTTAGG |
| **RM RPP30 Reverse 2** | SIV IPDA | ACATTCATGCCACTGCACTC |
| **RM RPP30 Probe 2** | SIV IPDA | HEX-ACAGGGTCTCACTTTGTTGTCCA-3IABkFQ |

### Supplementary Table 5: Cell annotation

| | CELL TYPE MARKERS | CELL TYPE |
|---|---|---|
| **CLUSTER 0** | CCR7, LEF1, CD3D,CD3E, CD4, CD8A, IL7R, TCF7, SELL, CD79A-, CD79B-, CD74- | **Naïve T cells** |
| **CLUSTER 1** | IGHM, CD79A, CD79B, CD74, MS4A1, FCRL2, BANK1, FCRLA, CD3D-, CD3E-, CD8A- | **B cells** |
| **CLUSTER 2** | CD8A-, CD3D, CD3E, CD4, CD79A-, CD79B-, CD74- | **CD4+ T cells** |
| **CLUSTER 3** | CD8A, CD3D, CD3E, NKG7, GZMK, GZMM, GZMB, GZMA, KLRD1, CD4-, CD79A-, CD79B-, CD74- | **CD8+ T cells** |
| **CLUSTER 4** | - | **-** |

**Note:** Resolution 0.1

**Supplementary Table 6: Cell annotation**

| | CELL TYPE MARKERS | CELL TYPE |
|---|---|---|
| **CLUSTER 0** | CCR7, LEF1, CD3D, CD4, CD8A-, RABGAP1L-, ITGA4-, ELMO1-, CD74-, CD79A-, CD79B- | **Naïve CD4+ T cells 1** |
| **CLUSTER 1** | CCR7, LEF1, CD3D, CD4, CD8A-, RABGAP1L-, ITGA4-, ELMO1-, CD74-, CD79A-, CD79B- | **Naïve CD4+ T cells 2** |
| **CLUSTER 2** | CD8A-, CD3D, CD4, IL7R, CCL5-, GZMK-, CD79A-, CD79B-, CD74- | **Memory CD4+ T cells 1** |
| **CLUSTER 3** | CD3D, CD3E, CD4, CD8-, CD79A-, CD79B-, SKAP1, LCK, PTPRC, IL7R, TRAC, LTB, CD2, CCL5-, GZMK-, KLRB1- | **Central memory CD4+ T cells** |
| **CLUSTER 4** | CD74, IGHM, DERL3, CD79A, CD79B, CD3E-,CD3D-, MS4A1, FCRL2, BANK1, FCRLA, RALGPS2, AIM2-, TCL1A, CD37, CD22 | **Proliferating B cells 1** |
| **CLUSTER 5** | CD8A-, CD3D, CD4, IL7R, CCL5-, GZMK-, CD79A-, CD79B-, CD74- | **Memory CD4+ T Cell 2** |
| **CLUSTER 7** | CD74, IGHM, DERL3, CD79A, CD79B, CD3E-,CD3D-, MS4A1, FCRL2, BANK1, FCRLA, RALGPS2, AIM2, VPREB3 | **Memory B cells 1** |
| **CLUSTER 8** | CD74, IGHM, DERL3, CD79A, CD79B, CD3E-,CD3D-, MS4A1, FCRL2, BANK1, FCRLA, RALGPS2, TCL1A, CD37, CD22, | **Proliferating B cells 2** |
| **CLUSTER 9** | CD74, IGHM, DERL3, CD79A, CD79B, CD3E-,CD3D-, MS4A1, FCRL2, BANK1, FCRLA, RALGPS2, AIM2, VPREB3 | **Memory B cells 2** |
| **CLUSTER 10** | GZMK, GZMM, GZMB, GZMA, LAG3, CCL5, TOX, PRF1, CD3D, CD3E, CD3G, CD8A, CD4-, CD74-, CD79A-, CD79B- | **Exhausted CD8+ T cells** |
| **CLUSTER 11** | CCL5, GZMB, GZMM, GZMK, CD3D, CD3E, CD3G, CD8A, CD4-, CD74-, CD79A-, CD79B- | **Cytotoxic CD8+ T cells** |
| **CLUSTER 12** | IL7R, TCF7, LEF1, SELL, CD3D, CD3E, CD8A, CD79A-, CD79B-, CD4-, CD74- | **Naïve CD8+ T cells 1** |
| **CLUSTER 14** | IL7R, TCF7, LEF1, SELL, CD3D, CD3E, CD8A, CD79A-, CD79B-, CD4-, CD74- | **Naïve CD8+ T cells 2** |
| **CLUSTER 15** | CD8A-, CD3D, CD4, CD79A-, CD79B-, CD74- | **CD4+ T cell subset** |

| | | |
|---|---|---|
| **CLUSTER 17** | CD8A-, CD3D, CD4, CD79A-, CD79B-, CD74-, CXCR5 | **Follicular CD4+ T cells** |
| **CLUSTER 18** | CD8A-, CD3D, CD4, IL7R, CCL5-, GZMK-, CD79A-, CD79B-, CD74- | **Memory CD4+ T Cell 3** |
| **CLUSTER 19** | CD74, IGHM, DERL3, CD79A, CD79B, CD3E-,CD3D-, MS4A1, FCRL2, BANK1, FCRLA, RALGPS2, AIM2, VPREB3, SSPN | **Memory B cells 3** |
| **CLUSTER 21** | FOXP3, TNFRSF18, IL2RA, TIGIT, CTLA4, IKZF2 | **Regulatory T cells** |
| **CLUSTER 23** | FCRL5, IGHM, CD79A, CD79B, CD74, CD3D-, CD3E- | **Unswitched memory B cells** |

**Note:** Resolution 2.0
